# Chromatin topology and the timing of enhancer function at the *hoxd* locus

**DOI:** 10.1101/2020.07.12.199109

**Authors:** Eddie Rodríguez-Carballo, Lucille Lopez-Delisle, Andréa Willemin, Leonardo Beccari, Sandra Gitto, Bénédicte Mascrez, Denis Duboule

## Abstract

The *HoxD* gene cluster is critical for proper limb formation in tetrapods. In the emerging limb buds, different sub-groups of *Hoxd* genes respond first to a proximal regulatory signal, then to a distal signal that organizes digits. These two regulations are exclusive from one another and emanate from two distinct TADs flanking *HoxD*, both containing a range of appropriate enhancer sequences. The telomeric TAD (T-DOM) contains several enhancers active in presumptive forearm cells and is divided into two sub-TADs separated by a CTCF-rich boundary, which defines two regulatory sub-modules. To understand the importance of this particular regulatory topology to control *Hoxd* gene transcription in time and space, we either deleted or inverted this sub-TAD boundary, eliminated the CTCF binding sites or inverted the entire T-DOM to exchange the respective positions of the two sub-TADs. The effects of such perturbations on the transcriptional regulation of *Hoxd* genes illustrate the requirement of this regulatory topology for the precise timing of gene activation. However, the spatial distribution of transcripts was eventually resumed, showing that the presence of enhancers sequences, rather than either their exact topology or a particular chromatin architecture, is the key factor. We also show that the affinity of enhancers to find their natural target genes can overcome the presence of both a strong TAD border and an unfavourable orientation of CTCF sites.

**SIGNIFICANCE STATEMENT:** Many genes important for vertebrate development are surrounded by series of remote enhancer sequences. Such regulatory landscapes and their target genes are usually located within the same chromatin domains, which appears to constrain the action of these regulatory sequences and hence to facilitate enhancer-promoter recognition and gene expression. We used the *HoxD* locus to assess the impact of modifying the regulatory topology upon gene activation in space and time. A series of chromosomal re-arrangements involving deletions and inversions reveals that the enhancer topology plays a role in the timing of gene activation. However, gene expression was often recovered, subsequently, illustrating the intrinsic capacity of some enhancers to find their target promoters despite an apparently adverse chromatin topology.

## INTRODUCTION

During embryonic development, the precise control of gene activation in both time and space largely relies on the activity of *cis*-regulatory sequences. Such regulatory elements include insulators, enhancers and repressive sequences that are either located in close proximity to the target gene or further away (Long et al., 2016). Large regulatory distances can be overcome by the three-dimensional organization of chromatin that takes place at different levels (Schoenfelder and Fraser, 2019). In this context, topologically associating domains (TADs) were defined as genomic intervals where chromatin interactions tend to take place more frequently than with adjacent regions (Dixon et al., 2012; Nora et al., 2012) and such domains are frequently understood as functional units that host enhancers and their target promoters (Furlong and Levine, 2018). Indeed, some key developmental genes are found under the control of regulatory domains that are contained within TADs, which harbour tissue-specific regulatory sequences or multiple acting enhancers that confer robustness and resilience (Amândio et al., 2020; Montavon et al., 2011; Osterwalder et al., 2018; Sagai et al., 2009; Will et al., 2017).

TADs and chromatin loops are thought to result from a loop extrusion mechanism that relies on the loading of the Cohesin multiprotein ring. This protein complex allows the extrusion of the chromatin fibre until it is stopped or retained by CTCFs bound with convergent orientations, or by the stalling of two forming loops (Fudenberg et al., 2016). The precise role(s) of these architectural proteins in gene expression has not yet been completely elucidated and genome-wide depletion either of CTCF, or of the cohesin complex, did not have a pervasive effect on gene expression levels and changes were clearly observed only at some genomic loci (Nora et al., 2017; Rao et al., 2017; Schwarzer et al., 2017; Soshnikova et al., 2010). Altogether, the relationship between chromatin topology and gene expression seems to be context-dependent and locus-specific. In some cases indeed, deleting CTCF binding sites led to an alteration of the genes nearby (Cuartero et al., 2018; Dowen et al., 2014; Hanssen et al., 2017; Hnisz et al., 2016), whereas other studies failed to reveal any obvious effects (de Wit et al., 2015). Most of these studies nevertheless did not monitor gene expression using time course protocols in a physiological situation *in embryo*.

*Hox* clusters have been used as a paradigm of long-range regulation. The *HoxD* cluster It is localized between two large TADs and acts itself as a boundary region, due to the high concentration of CTCF binding sites and their opposed orientations (Andrey et al., 2013; Rodríguez-Carballo et al., 2017). The centromeric domain (C-DOM) controls *Hoxd* genes transcription during the late, second phase of limb development, which accompanies the emergence of digits (Lonfat et al., 2014; Montavon et al., 2011). The telomeric domain (T-DOM) controls the early phase of transcription in limb buds (Andrey et al., 2013; Tarchini and Duboule, 2006; Zakany et al., 2004), as well as in the caecum (Delpretti et al., 2013) and the mammary buds (Schep et al., 2016). T-DOM is divided into two sub-TADs by a chromatin boundary (CS38-40) containing three bound CTCFs, all oriented towards the *HoxD* cluster (Rodríguez-Carballo et al., 2017) (Fig. 1A)

**Figure 1.**
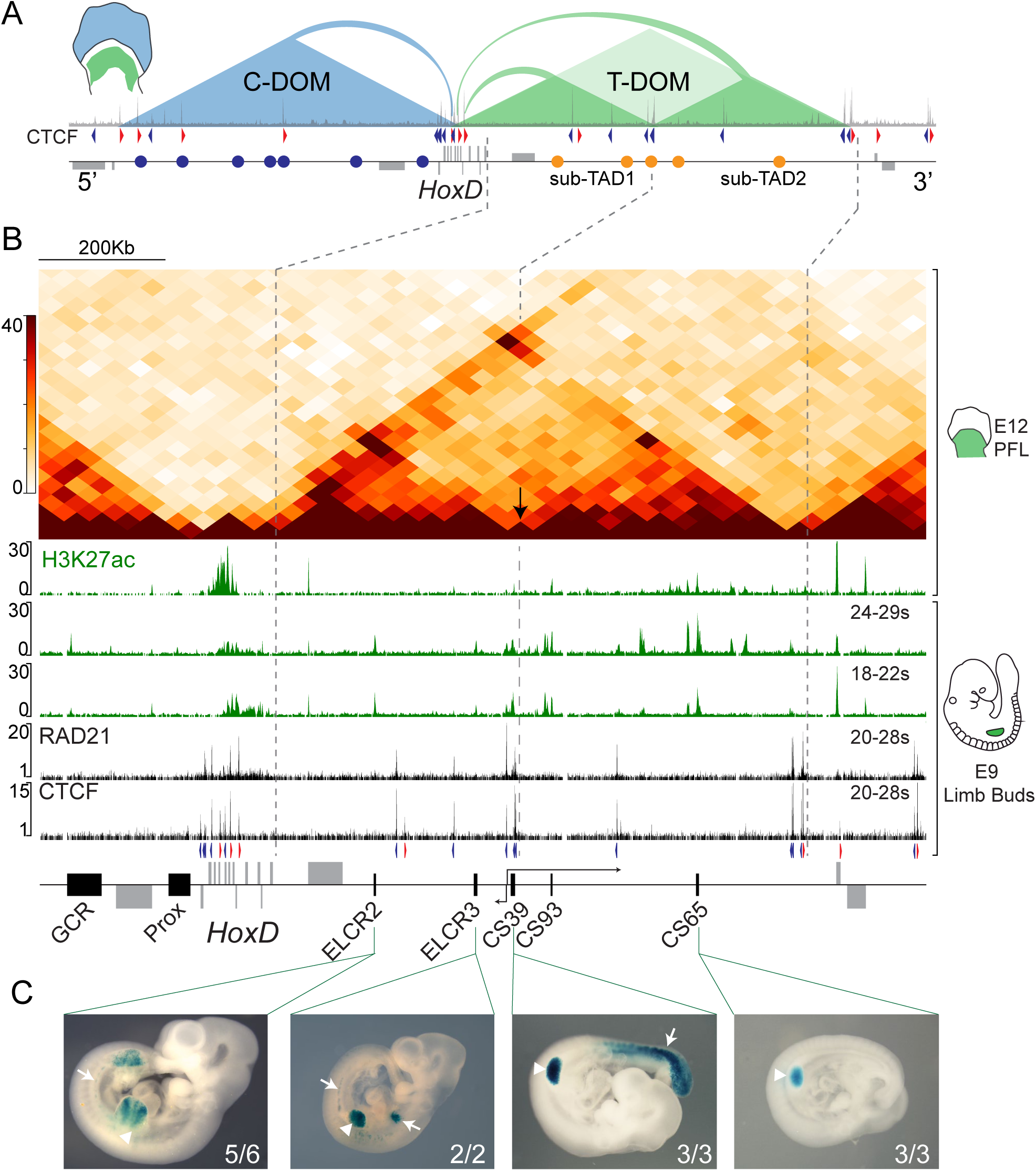
The *HoxD* locus and its regulatory landscapes. **A**. Scheme of the *HoxD* locus showing the gene cluster surrounded by limb enhancers (blue and yellow circles) and neighbouring genes (grey boxes). A ChIP of CTCF in E12.5 proximal forelimbs is shown, as well as the orientation of their binding sites (blue and red arrowheads). The regulatory domains C-DOM (blue) and T-DOM (green, split in two sub-TADs) are represented as triangles. On the left, a E12.5 limb diagram shows the tissue where C-DOM and T-DOM are active, respectively. **B**. Magnification of *HoxD* and the T-DOM region. Top panel shows a Hi-C map of E12.5 proximal limb (data from (Rodríguez-Carballo et al., 2017)). Arrow shows the sub-TAD division. The green tracks are ChIP datasets of H3K27ac in E12.5 proximal limb and E9 forelimb buds at 24 to 29 and 18 to 22 somite stage. The black tracks (bottom) are ChIP tracks of RAD21 and CTCF of E9 forelimbs at 20 to 28 somite stage. **C**. *LacZ* staining of mouse embryos showing limb (arrowheads) and mesodermal enhancer (arrows) activity of several transgenic constructs at E10.5 (ELCR2) and E.9.5 (ELCR3, CS39 and CS65).

In the early limb bud, T-DOM is activated at E9.0, leading to the first wave of *HoxD* colinear transcription, coinciding with the establishment of chromatin interactions between the newly activated genes (*Hoxd9-Hoxd11*) and part of T-DOM (Andrey et al., 2013). At E12.5, cells transcribing these genes are found in the proximal part of the limb buds, which will generate the arm and the forearm. At this stage, the distribution of interactions with T-DOM shows a clear topological segregation, with 3’-located genes (*Hoxd1* to *Hoxd8)* interacting mostly with the first sub-TAD, whereas 5’-located genes (*Hoxd9* to *Hoxd11*) associate in priority with the more distant sub-TAD (Andrey et al., 2013; Rodríguez-Carballo et al., 2017), suggesting a functional compartmentalisation of T-DOM. All limb-specific enhancers were thus far associated to the distant sub-TAD, starting at the sub-TAD boundary and extending up to *Hnrnpa3*, including the CS65 and CS93 enhancers (Andrey et al., 2013; Yakushiji-Kaminatsui et al., 2018).

In this work, we set up to assess whether a correlation exists between the precise temporal and spatial transcriptional activation of *Hoxd* genes in proximal limbs on the one hand, and a fine topological organisation of its regulatory landscape, on the other hand, or whether the mere presence of series of enhancers within T-DOM is necessary for *HoxD* regulation, regardless of their intrinsic organisation. We show that while the overall chromatin architecture determines the correct timing of gene activation, enhancer-promoter communication can be successfully established along with limb bud development, even after the engineering of major topological modifications, including the positioning of a strong TAD border in between them.

## RESULTS

### Multiple early limb enhancers in T-DOM

Hi-C profiles from several cell types have previously revealed that the *HoxD* cluster is positioned at the boundary between two TADs. T-DOM, i.e. the TAD located telomeric to the gene cluster, is necessary for the transcription of *Hoxd* genes both during limb budding and, subsequently, in the formation of the proximal segment of the prospective arm. Instead, the C-DOM controls *Hoxd* gene expression in developing digits, at later time points (Fig. 1A). From E9.5 to E12.5, T-DOM shows specific activation and decommissioning dynamics (Andrey et al., 2013), which correlates with its 3D conformation, as only the more distant T-DOM sub-TAD (Fig. 1A, sub-TAD2) remains active at late (E12.5) embryonic stages. Most limb enhancers described thus far are located within this chromatin domain, in particular the CS39, CS65 and CS93 sequences (Andrey et al., 2013; Beccari et al., 2016; Yakushiji-Kaminatsui et al., 2018).

In order to characterize the onset of activation of T-DOM at the earliest time of *Hoxd* gene transcription, in the incipient limb bud, we micro-dissected E9 forelimb buds and pooled them into two groups corresponding to embryos either between 18 to 22 somites (or early E9), or between 24 to 29 somites (or late E9). ChIP of H3K27ac, a histone mark associated with enhancer activity and gene expression, revealed that most of the acetylated regions were located in the second sub-TAD, which seems particularly active in 24 to 29 somites old limb buds (Fig. 1B). Two H3K27ac-positive regions were nevertheless identified in the first sub-TAD in E9 limb buds, which were not present in E12.5 proximal limb cells. In contrast to CS39, CS65 and CS93, however, these two early limb control regions (ELCR2 and ELCR3) were not found fully conserved in chicken, albeit they are present in all mammals (Fig. S1 and (Andrey et al., 2013; Yakushiji-Kaminatsui et al., 2018)). Transgenic analysis of both ELCR2 and ELCR3 showed strong *LacZ* expression in E9 limb buds, which coincides with the expression of CS39 and CS65 transgenes (Fig. 1C, arrowheads), as well as in other mesoderm derivatives (Fig. 1C, arrows).

To evaluate potential changes in the global architecture of T-DOM along with developmental timing, we looked at the binding profiles of both CTCF and the Cohesin subunit RAD21. The ChIP profiles of these architectural proteins using limb buds from 20 to 28 somites embryos did not substantially differ from the profiles obtained in E12.5 proximal limb (Fig. 1B and Fig. 2A, B in (Rodríguez-Carballo et al., 2017)). Most of the CTCF binding sites had a convergent orientation in relation to the *HoxD* cluster, including the three bound CTCFs found within the CS38-40 region, the boundary region that divides T-DOM into its two sub-TADs (Fig. 1B, arrow).

**Figure 2.**
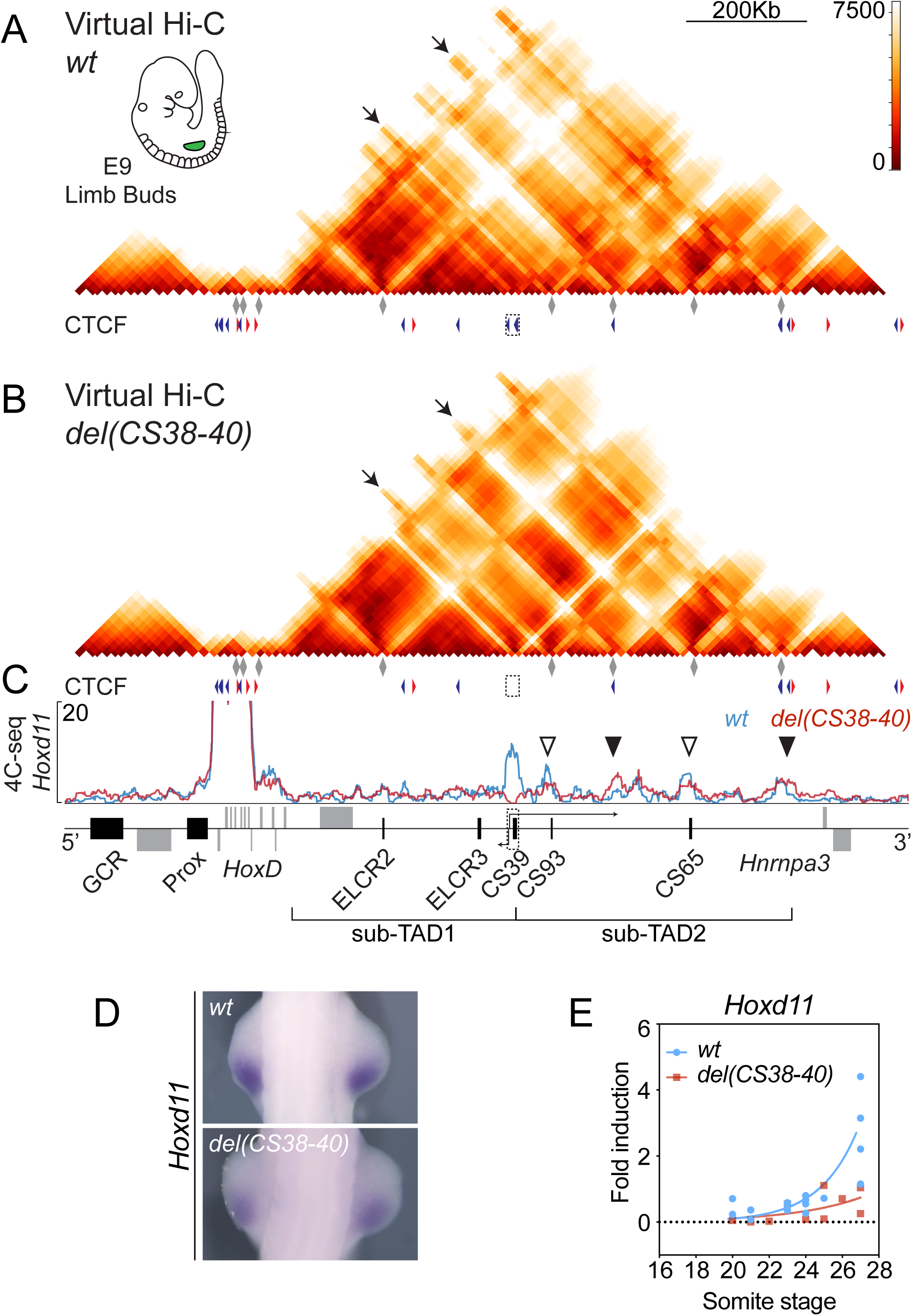
Deletion of region CS38-40. **A, B**. Virtual Hi-C maps of wild-type (**A**) and *del(CS38-40)* (**B**) as reconstituted from 4C-seq datasets of E9 forelimb buds. The 4C viewpoints used to compose the matrices (grey diamonds) as well as the positions and orientations of the CTCF sites are depicted below. The scales on the right represent the computed distances between the different bins. **C**. 4C-seq profiles of *Hoxd11* of E9 of *wt* and mutant forelimb buds (blue and red lines, respectively). The deleted region is shown as a dashed box around CS39, from which two non-coding transcripts are generated in the wild-type. Arrows arising from the dashed box around CS39 represent the *Hog* and *Tog* lncRNAs in the wild-type. **D**. WISH analysis of *Hoxd11* in E9 forelimbs (approximately 26 to 27 somites). **E**. Comparison of individual RT-qPCR values (*wt* n=17 and mutant n=11) of *Hoxd11* in E9 forelimb buds at different somite stages. An exponential fit is represented out of the real RT-qPCR values.

### Deletion of the T-DOM sub-TAD boundary

We asked whether such a partitioning of T-DOM into two subdomains was mandatory for this early limb bud regulation to be properly implemented. We to merged both domains by deleting the CS38-40 region (Fig. S2A), which contains three CTCFs binding sites as well as the CS39 limb enhancer and the transcription start site (TSS) of the *Hog* and *Tog* lncRNAs (Delpretti et al., 2013; Rodríguez-Carballo et al., 2017), which approximatively coincides with a CpG island. We performed 4C-seq experiments in E9.5 mutant forelimb buds using several viewpoints distributed both along T-DOM and inside the *HoxD* cluster. We then compiled all data from the different viewpoints into a virtual Hi-C matrix using the 4Cin software (Irastorza-Azcarate et al., 2018), which we adapted to plot the relative distances in a linear manner according to the real genomic coordinates. Because the 4Cin processing has an inherent variability that leads to the generation of different models, we assessed the correlation between twenty iterations and clustered them, thus displaying a merged average (see materials and methods).

When applied to control limbs, the 4Cin approach generated a map of computed distances at T-DOM that resembled the expected contact distribution of a wild-type Hi-C matrix, including the subdivision of the domain in two sub-TADs, as well as specific contacts between CTCF-bound regions and enhancer-promoter interactions (Fig. 2A). Using the same viewpoints, we confirmed that the deletion of region CS38-40 affected the spatial organization of this regulatory domain. We observed a substantial increase in the interactions established between the two sub-TADs, leading to their fusion into a single domain (Fig. 2B). The increase in contacts between the two sub-TADs could be observed when comparing any of the derived cluster representations (Fig. S2B, C). Also, the interactions established by the *HoxD* cluster throughout the regulatory domain decreased (Fig2. A, B, arrows at CS93 and CS65), even though they could still be identified.

More specifically, we analysed the interaction profile of the *Hoxd11* gene, whose expression in the posterior part of the E9.5 developing limb bud is maintained until E12.5, in the proximal limb. 4C-seq data revealed that in wild-type E9.5 limb buds, *Hoxd11* strongly interacted with both region CS38-40 and the more distant sub-TAD (Fig. 2C). Upon deletion of the sub-TAD border, a modest increase in interactions was detected in the bound CTCF sites located 3’ to region CS38-40 and at the telomeric TAD border close to *Hnrnpa3* (Fig. 2C, bold arrowheads). However, the contacts did not increase substantially along the region initially corresponding to the second sub-TAD. Instead, interactions were reduced with the CS93 and CS65 limb enhancers (Fig. 2C, open arrowheads).

We assessed whether these alterations in contact distribution translated into changes in gene expression pattern. Whole mount RNA in situ hybridization (WISH) showed a slight but visible decrease in *Hoxd11* expression at E9.5 (Fig. 2D). To verify this observation, we performed RT-qPCR on forelimb buds dissected from embryos aged between 20 and 28 somites and plotted their individual values (Fig. 2E). The dynamics of *Hoxd11* expression in control forelimb buds followed a strong increase right after the 24 somites stage. On the contrary, this dynamics in *Hoxd11* mRNA was not observed in the mutant limb buds where the increase was not as fast (Fig. 2E). This was further confirmed by RNA-seq experiments showing that *Hoxd10* and *Hoxd12* had a delayed onset of transcription, while more anterior genes (i.e. *Hoxd4*) did not seem to be affected at all (Fig. S2D). These effects could either be a consequence of the distinct spatial reorganization of T-DOM or be due to the removal of CS39 enhancer.

To explore these possibilities, we used a CRISPR/Cas9 approach to eliminate the binding of CTCF to the three motives positioned within region CS38-40. We initially deleted 26 bp of the CTCF binding site located in region CS38 (*delCTCF(CS38)*; Fig. S3A), preserving both the neighbouring CpG island and the TSS of the *Hog* and *Tog* lncRNAs (Fig. S2A). On top of this first intervention, we generated a 1.5 kb large deletion that removed the two binding sites located around CS40, without removing the H3K27ac-enriched region localised around CS39 (*delCTCFs(CS38;CS40)*; Fig. S2A, S3A). We confirmed by ChIP that CTCF binding was no longer detected at any of these locations or elsewhere in this short DNA interval (Fig. S2A). The deletion of the three CTCF sites led to a merge of the sub-TADs at E12.5 (FigS3B, C), thus confirming the importance of these bound proteins in the establishment of this specific topological structure. We analysed gene expression and observed that *Hoxd11* was briefly delayed in its activation, a lag that was rapidly resumed to generate a late pattern indistinguishable from wildtype (Fig. S3D). These results indicated that the merging of the two T-DOM sub-TADs moderately affected the onset of *Hoxd* gene expression in early limb buds, with a stronger effect observed in the absence of the CS39 enhancer.

### Reinforcing a sub-TAD separation

We next set up to engineer the opposite situation, i.e. to produce a more robust separation between the two sub-TADs such as to isolate them from one another as bona fide TADs. This was achieved by generating an inversion of the region comprising the three CTCF binding sites (the *inv(CS38-40)* allele). In this configuration, the three CTCF sites now converged towards the strong telomeric TAD border at the 3’ end of the domain. In this allele, the three CTCF sites were still occupied, as expected (Fig. S4A). A virtual Hi-C pattern of limb bud cells dissected from this mutant stock revealed that, as expected, the inversion of this region had strengthened the segregation of the two sub-TADs. Concomitantly, it also strongly reduced the general contacts between the *HoxD* cluster and the CS93 and CS65 regions (Fig. 3A, B; arrows). This was illustrated by using *Hoxd11* as a 4C-seq viewpoint, showing a reduction of interactions over regions CS38-40, CS93 and CS65 (Fig. 3C), similar to what had been noted in the *del(CS38-40)* allele (Fig. 2C). These topological changes also correlated with a delay of *Hoxd11* expression (Fig. 3D, E), which was stronger than in the CTCF mutant alleles. This delay was nevertheless not pervasive, for it did not affect all *Hoxd* mRNAs equally. For example, *Hoxd9* did not show a clear transcriptional decrease, even at early stages (Fig. S4B), whereas more ‘posterior’ genes (like *Hoxd11*) seemed to be more affected. All these delays, however, were subsequently rescued and, in E12.5 forelimb buds, changes in expression patterns could hardly be scored when comparing the *inv(CS38-40)* allele to wild-type littermates (Fig. S4C).

**Figure 3.**
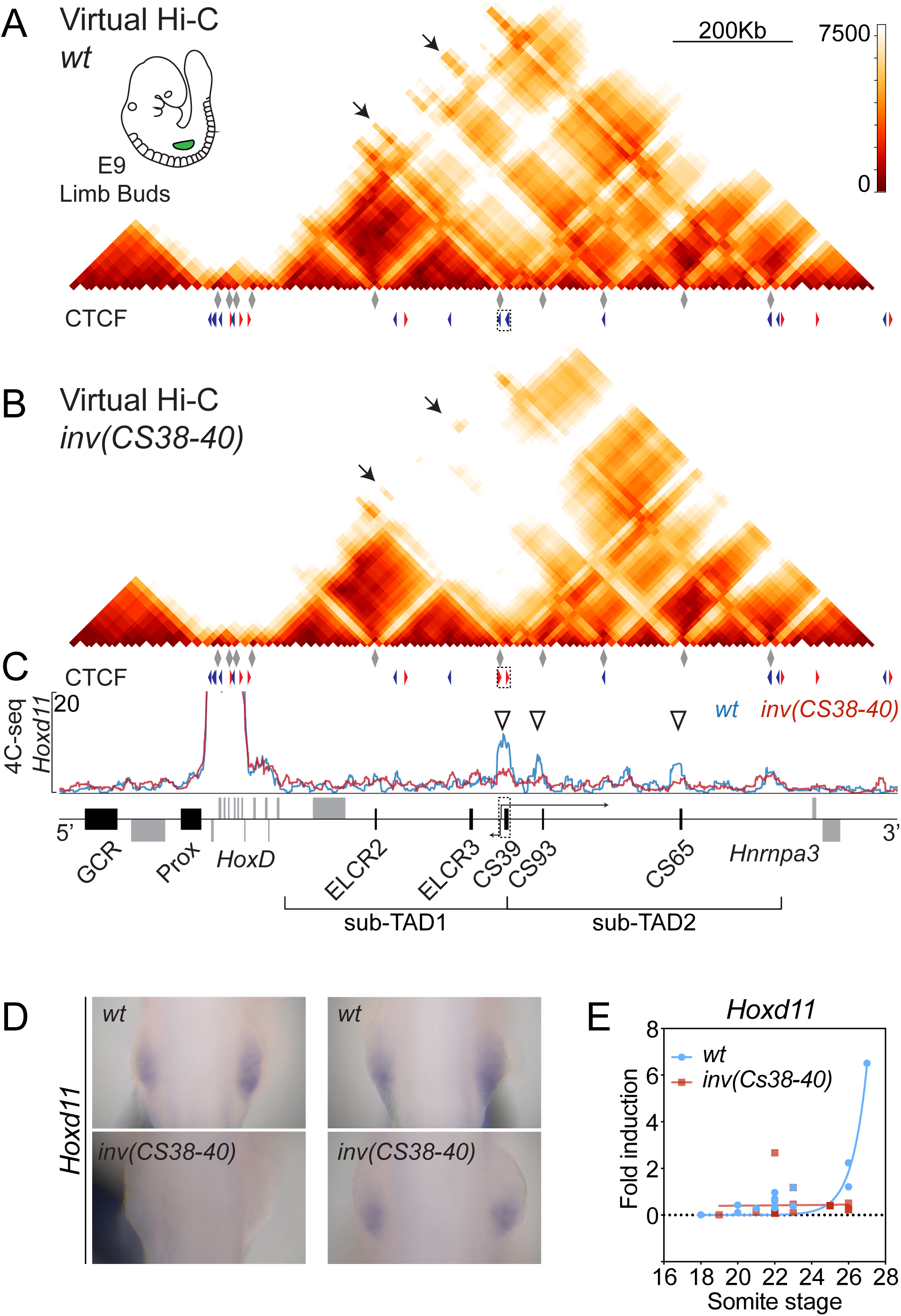
Inversion of region CS38-40. **A, B**. Virtual Hi-C maps of wild-type (**A**) and the *inv(CS38-40)* allele (**B**) from E9 forelimb buds, as reconstituted from several 4C-seq viewpoints (grey diamonds). The triangles showing CTCF orientation of region CS38-40 are inverted in the mutant. **C**. 4C-seq profiles of *Hoxd11* for *wt* and mutant E9 forelimb buds (blue and red lines, respectively). The inverted region is shown as a dashed box around CS39. Empty arrowheads show areas of decreased interaction in the *inv(CS38-40)* allele. **D**. WISH analysis of *Hoxd11* in E9 (18 to 22 somites) forelimb buds (earlier time point on the left). **E**. Comparison of RT-qPCR values (*wt* n=16 and mutant n=17) of *Hoxd11* in E9 forelimb buds at different somite stages. An exponential fit is represented out of the real RT-qPCR values.

### Invertion of T-DOM

Altogether, these genomic alterations did not produce long-lasting effects upon the transcription of *Hoxd* genes in limb buds. One possibility is that T-DOM contains several other CTCF binding sites, most of them displaying an orientation convergent to those numerous sites present in the telomeric part of the *HoxD* cluster itself. In this context, it is possible that such CTCF sites within T-DOM may assist remote enhancers reaching targets, regardless of small rearrangement occurring at their vicinity. We thus set up to invert the entire T-DOM such as to produce two distinct inversion alleles, one containing a strong TAD border between the inverted T-DOM and the *HoxD* cluster, and the other one lacking this TAD border (Figs. 4 and 5, Bd).

**Figure 4.**
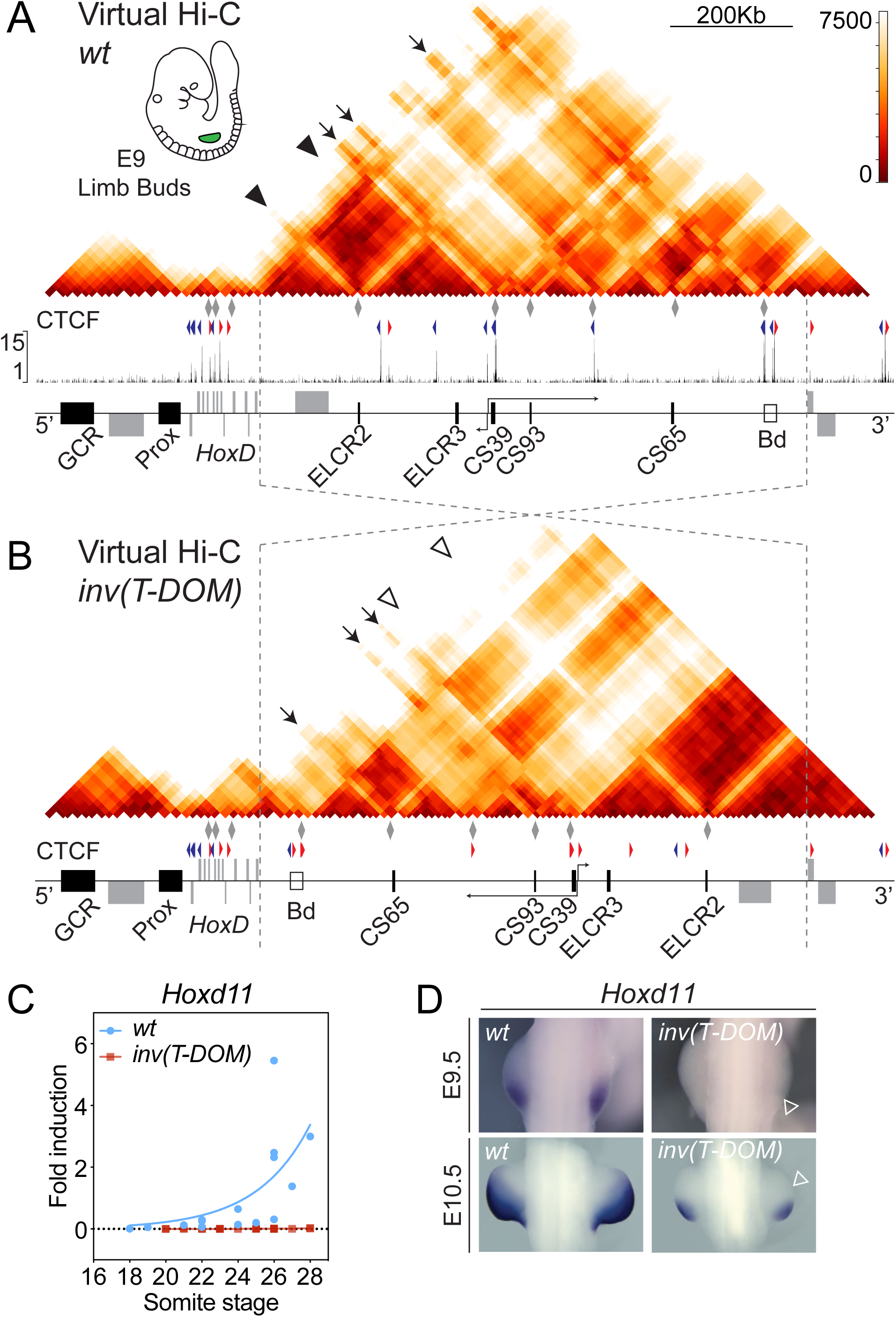
Inversion of T-DOM in the presence of a TAD border. **A, B**. Virtual Hi-C maps of wild-type (**A**) and *inv(T-DOM)* mutant mice (**B**) from E9 forelimb buds, as reconstituted from several 4C-seq viewpoints (grey diamonds). Dashed lines indicate the inverted region. The distant boundary is marked as an empty box (Bd) at the end of T-DOM in the wild-type and close to the *HoxD* cluster in the mutant allele. The CTCF orientations in the *inv(T-DOM)* allele are inverted accordingly. In the mutant allele, empty arrowheads represent lost interactions that can be scored in the *wt* (bold arrowheads). **C**. RT-qPCR values (*wt* n=17 and mutant allele n=21) of *Hoxd11* in E9 forelimb buds at different somite stages. An exponential fit is represented out of the real RT-qPCR values. **D**. WISH images of forelimb buds at E9.5 (top, approximately 20 somites) and E10.5 (bottom) of *wt* and the *inv(T-DOM)* mutant allele. The loss of expression in the mutant is indicated as empty arrowheads.

**Figure 5.**
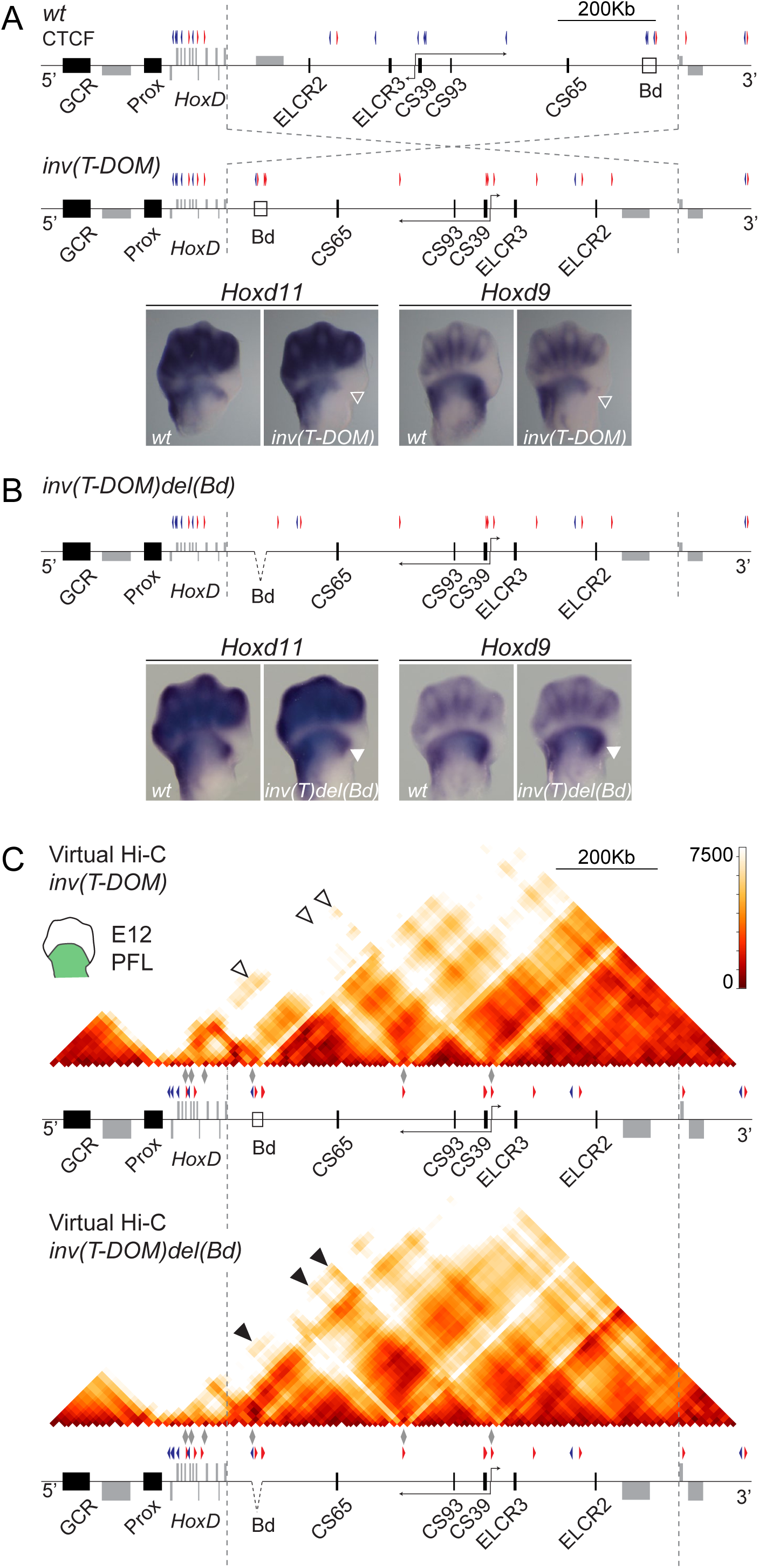
Inversion of T-DOM in the absence of a TAD border. **A**. Scheme of the re-arranged *HoxD* locus after inversion of T-DOM. The orientation of CTCF sites are represented. WISH images of *Hoxd11* and *Hoxd9* expression pattern in E12.5 forelimbs are shown for both *wt* and *inv(T-DOM)* mutant embryos. An empty arrowhead demarcates the loss of expression in the anterior part of the proximal limb domain for both *Hoxd9* and *Hoxd11*. **B**. Scheme showing the deletion of the TAD boundary (Bd) on top of the *inv(T-DOM)* allele. WISH of *Hoxd11* and *Hoxd9* using E12.5 *wt* and *inv(T-DOM)del(Bd)* -*inv(T)del(Bd)*-mutant forelimbs show the same patterns with the re-apparition of the anterior domain missing in the *inv(T-DOM)* allele (arrowhead). **C**. Virtual Hi-C maps of E12 proximal limb buds using either *inv(T-DOM)*, or *inv(T-DOM)del(Bd)* mutant limb bud cells. Both matrices were mapped on the artificial *inv(T-DOM)* genome. Bold arrowheads indicate the increase in contacts with enhancers CS65, CS93 and CS39 (indicated with empty arrowheads in *inv(T-DOM)*) after the deletion of the TAD boundary (Bd).

We induced an inversion of the entire T-DOM by targeting CRISPR guides at both sides of this large regulatory domain. To be as inclusive as possible, the breakpoints of this inversion were selected close to the 3’ end of the *Hoxd1* gene and 5 kb upstream the TSS of the *Hnrnpa3* gene, respectively (Fig. 4A, B, dashed lines). Due to the position of the latter breakpoint, the *Hnrnpa3* TAD border (Fig. 4A, Bd) was inverted along with T-DOM and placed just between the *HoxD* cluster and the inverted T-DOM. Upon inversion, a substantial loss of contacts was scored by using 4Cin on E9.5 limb bud cells, all along the T-DOM and particularly in the region containing the ELCR2 and ELCR3 enhancers, which had been located further away (Fig. 4A, B, black and open arrowheads).

In contrast, the CS65, CS93 and CS39 enhancers, which were initially located further telomeric, were now re-positioned much closer to the *HoxD* cluster in the inverted allele. These enhancers were nevertheless separated from their natural *Hoxd* target genes by a strong and very efficient TAD border (Fig. 4B, Bd). Noteworthy, however, clear interactions could still be observed between *Hoxd* genes and these regulatory regions, although clearly diminished when compared to the control (Fig. 4A, B, compare arrows). These interactions took place in spite of this TAD border (Bd), which otherwise tightly isolated the inverted T-DOM from the *HoxD* cluster, creating a new and well identified TAD. Again, these changes were observed regardless of the model that was generated in our virtual Hi-C approach (Fig. S5A, B). While the interactions between the *HoxD* cluster and the T-DOM enhancers were remarkably weaker, they were observed, in particular when looking at the 4C-seq profile of *Hoxd11* (Fig. S5C).

The inversion of the whole regulatory domain was accompanied by a severe delay in the onset of *Hoxd11* expression, which was not detectable before E10.5 at the most posterior aspect of the growing bud (Fig. 4C, D), likely due to the weak interactions with T-DOM enhancers. This strong variation in the timing of expression was subsequently translated into an absence of both *Hoxd9* and *Hoxd11* transcripts in the most anterior part of the proximal expression domain at E12.5, i.e. when the transcript domains have reached their final spatial deployments (Fig. 5A; white arrowheads). Altogether, these results demonstrated that, in this case, a delay in *Hox* genes activation impacted upon the spatial distribution of their transcripts. They also illustrate that enhancers seem to be able to still contact their natural target genes, despite the presence of a strong ectopic TAD border in between.

### Enhancer tropism over chromatin topology

To see whether the altered expression timing and patterns of *Hoxd9* and *Hoxd11* were due either to the mere inversion of T-DOM, or to the introduction of a strong new TAD border between the gene cluster and T-DOM, we further deleted the boundary region on top of the inverted allele to produce the *inv(T-DOM)del(Bd)* mutant line (Fig. 5). The deleted 20 kb large boundary region, which normally tightly isolates T-DOM from its more telomeric TAD, contained three CTCF binding sites. Fetuses carrying this additional deletion fully recovered wild type expression patterns for both *Hoxd9* and *Hoxd11*, with expression domains in the limb buds undistinguishable from their wild-type counterparts (Fig. 5B). In particular, the proximal-anterior transcript domain lacking in *inv(T-DOM)* embryos (Fig. 5A, open arrowheads) was fully rescued after deletion of the ectopic TAD border (Fig. 5B, bold arrowheads).

This recovery in expression was concomitant to a clear increase in interactions between the *HoxD* cluster and various T-DOM limb enhancers when looking both at the virtual Hi-C matrices (Figs. 5C and S6A) and to the *Hoxd11* 4C-seq profile (Fig. S6B). The re-establishment in the spatial deployment of transcripts at day E12.5 was however not observed at the earliest stages analysed, which still showed an important time lag in target gene activation, even though the ectopic TAD border had been removed (Fig. S6C). These results showed that the mere inversion of the regulatory domain had an impact upon the onset of *HoxD* expression. The observed delay could nevertheless be caught up in a few days, a recovery that was not completely possible when the telomeric TAD border was present between the enhancers and the target *Hoxd* genes.

## DISCUSSION

The fine-tuned regulation of genes involved in developmental processes is often achieved by complex regulatory landscapes, which can extend up to megabases around the target gene(s). Such regulatory landscapes generally match the extents of TADs and contain all the enhancers necessary for the various expression specificities. Even though a clear causal relationship is difficult to establish, the prevalent model is that TADs somehow restrict the sphere of operation for such regulations, by providing a spatial unit where genes can be properly controlled, in isolation from their neighbours. The action of enhancers is thought to depend on their 3D spatial proximity to the target promoters they regulate, a hypothesis supported by several lines of evidence (reviewed in (Long et al., 2016; Schoenfelder and Fraser, 2019)). Recent reports, however, have challenged this view showing that transcriptional activity does not always correlate with a direct promoter-enhancer physical interaction (Alexander et al., 2019; Benabdallah et al., 2019; Fukaya et al., 2016).

In this work, we used the *HoxD* locus and one of its two flanking regulatory landscapes as a paradigm to look at the effect of modifying the regulatory topology upon the precisely orchestrated transcription of this series of genes. We engineered several rearrangements within the regulatory domain to determine the impact of both the distribution of enhancer sequences, the presence and orientation of CTCF binding sites and the ectopic introduction of a TAD border between the promoters and the corresponding enhancers. We conclude that, while the global TAD architecture may serve to properly implement the regulatory modalities in time, major re-arrangements do not critically modify the regulatory outcome at a later stage, making enhancer-promoter contacts very resilient and somewhat poorly dependent from the architectural context.

### A split regulatory landscape

T-DOM is normally divided into two sub-TADs at the level of region CS38-40 (Andrey et al., 2013), a region that contains three CTCF sites with an orientation convergent to that of numerous sites within the *HoxD* cluster (Rodríguez-Carballo et al., 2017). The deletion of this border region expectedly led to the fusion of the two sub-TADs. However, rather than re-enforcing contacts between enhancers and promoters in the *de novo* created single TAD, enhancer-promoter contacts tended to decrease. Therefore, the presence of these two sub-domains within T-DOM favours maximal efficiency in the regulatory outcome (see summary scheme in Fig. 6). One potential explanation is that it is not the global structure itself that is important but instead, the presence of three CTCF binding sites that may trigger part of the necessary interactions, in particular due to their shared orientation towards the *HoxD* cluster. In this deleted allele, *Hoxd* genes were expressed rather normally but with a clear delay in their activation.

**Figure 6.**
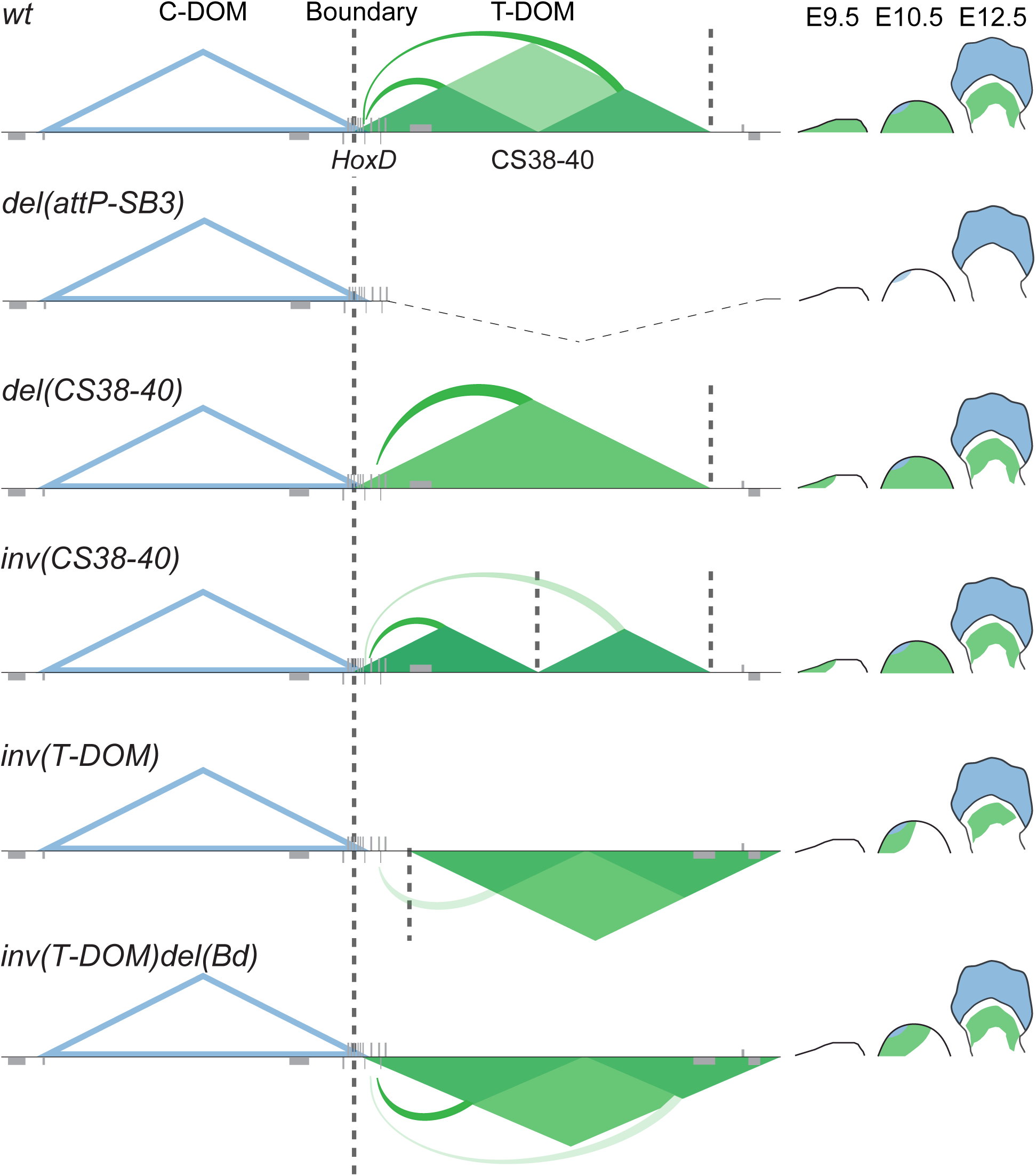
Summary of the regulatory effects observed in the various alleles. The C-DOM TAD (blue) is only active in distal cells during late limb development and hence it does not interfere with the current alleles. T-DOM (green) controls the transcriptional onset of *Hoxd* genes during early development (E9.5) and maintenance in the proximal limb bud until day 12.5. On top, the deletion of T-DOM abolished *HoxD* gene expression in the proximal limb (Andrey et al., 2013). Deletion of region CS38-40 leads to the merging of the two sub-TADs and a short delay in *Hoxd* genes activation. Inversion of the same region increases the segregation of the distant sub-TAD, while also delaying expression, which is subsequently rapidly recovered. The inversion of T-DOM and introduction of an ectopic TAD boundary hampers access of *Hoxd* genes to the domain. The delay in the activation is more severe. There is also an impact on the spatial distribution of transcripts in the anterior aspect of the proximal domain. The further deletion of this TAD border restores promoter-enhancer interactions at a later stage, whereas a strong impact is still visible during the earliest stages of limb development.

A more precise deletion strategy removing these three CTCF sites led to a similar fusion between the sub-domains. However, the effect upon *Hoxd* gene transcription was even milder that in the deletion of the boundary, likely because the full boundary deletion also included the CS39 enhancer, which was left in place in the CTCF deletion allele. In the latter case, mRNAs accumulation was also delayed, but even less that in the first allele. Therefore, it seems that the presence of these CTCF sites, rather than the global structure that they help to organize, as well as the full collection of limb enhancers are the key elements to properly activate the target genes in time.

The importance of CTCF sites and/or of their orientation for chromatin interactions was previously predicted *in silico* and illustrated experimentally at a variety of specific loci (Guo et al., 2015; Sanborn et al., 2015; de Wit et al., 2015). In developing tissues, the presence of bound CTCF in a specific locus favoured the ectopic action of some clustered enhancers when placed in a different TAD (Kraft et al., 2019). Here we show that when the entire boundary region containing the three sites was inverted, the isolation of the two sub-TADs became much stronger due to the convergence of these inverted sites with the natural telomeric 3’ TAD border, giving rise to two qualified TADs. Despite this accentuated split, which further isolated many limb enhancers from the target *Hoxd* genes, the transcriptional impact was once again restricted to the onset of their expression, similar to the effect of deleting the sub-TAD boundary. These results confirmed that the partition of T-DOM into sub-TADs may not respond to any particular regulatory necessity, at least in limb cells. Instead, it may be a consequence of CTCF being engaged into facilitating enhancer-promoter contacts.

### Enhancers topology and regulatory heterochronies

It is often argued that chromatin architecture is instrumental to ensure the proper temporal dynamics of gene activation (see e.g. (Furlong and Levine, 2018)). However, in the case where regulatory landscapes contain multiple enhancers with either identical (Hörnblad et al., 2020) or related (Montavon et al., 2011; Osterwalder et al., 2018) specificities, it is less clear whether the respective positions of these enhancers and their distances to one another are critical factors for target gene activation. T-DOM contains multiple limb bud enhancers over 800 kb, which tend to be distributed far from the gene cluster, interspersed with CTCF sites. The large engineered inversion of T-DOM lacking the TAD border gave us a rather clear answer to this question, at least regarding this particular locus. While *Hoxd* genes were importantly delayed in their activation at E9.5, their expression patterns at E12.5 were indistinguishable from control limb buds. This once more points to the separation between two distinct regulatory aspects; on the one hand, the full series of enhancers will end up delivering their integrated information, regardless of their global organisation within the landscape. On the other hand, an appropriate order and/or chromatin organisation will help to properly orchestrate this process.

While similar chromosomal rearrangements have been engineered at other developmental loci, it is difficult to propose a synthetic view of the results, for several parameters are usually involved and mixed with one another such as the presence or absence of TAD borders and/or CTCF sites, as well as the presence of enhancers, their relative distribution or their displacement related to their target genes. Deletions and duplications at the *Ihh* locus disrupted the communication with multiple enhancers, leading to limb malformations (Will et al., 2017). Also, re-arrangements of the TAD containing *Shh* and its enhancers led to deleterious effects on gene activation and concurrent phenotypes (Symmons et al., 2016). Yet, at this specific locus, moderate topological modifications did not elicit any severe limb defects, suggesting that enhancer-promoter communication may not rely only on a sustained 3D structure and that sporadic interactions may be sufficient (Alexander et al., 2019; Benabdallah et al., 2019; Fukaya et al., 2016). Expectedly, stronger phenotypes were obtained by deleting the ZRS enhancer region, as this sequence is the only known limb enhancer in this landscape (Lettice et al., 2017; Paliou et al., 2019; Williamson et al., 2019). The transitory effects observed when modifying T-DOM could either reflect a normal enhancer deployment delayed by changes in the 3D context or, alternatively, a novel organisation in enhancer-promoter interaction due to the known cooperative capacity that neighbouring enhancers can display during embryonic development (Amândio et al., 2020; Bolt and Duboule, 2020; Osterwalder et al., 2018). Similarly, the deletion of the TAD border and inversion of the regulatory domain at the *Sox9*/*Kcnj2* locus only had mild effects on gene expression (Despang et al., 2019).

### The resilience of enhancer-promoter interactions

In its initial form, the inversion of T-DOM introduced a strong ectopic TAD border between the enhancers and their target genes, in addition to the re-orientation of all CTCF binding sites. In this allele (Fig. 6), accordingly, the access of *Hoxd* genes to their cognate limb enhancers was dramatically reduced. While a severe delay was scored in transcriptional activation, some interactions surprisingly remained between the gene cluster and the regulatory domain, despite the latter being clearly in a distinct TAD. These contacts could even resume expression of *Hoxd* genes in proximal limb cells, although with a truncated spatial distribution. This observation is slightly at odds with the view of TAD borders restricting the access to neighbouring enhancers and delimiting regulatory interactions (van Bemmel et al., 2019; Franke et al., 2016; Lupiáñez et al., 2015; Nora et al., 2012; Rodríguez-Carballo et al., 2017). Here, despite the presence of a strong TAD border and the inversion of CTCF sites, which clearly led to the formation of a new TAD excluding the *HoxD* cluster, some enhancers-promoter interactions could still occur at a sufficient level to eventually produce detectable mRNAs in the expected proximal domain, thereby indicating that such contacts have intrinsic driving forces and do not entirely depend upon an instructive 3D context.

### Colinear regulation and phenotypic effects

During early limb bud development, *Hoxd* genes are activated in a time sequence that follows their respective positions along the gene cluster (Dollé et al., 1989; Tarchini and Duboule, 2006). The mechanism underlying this temporal colinearity process has been studied by intensive chromosome engineering whereby the order and/or presence of genes were modified, as well as their physical relationships with the adjacent regulatory landscapes (see references in (Tschopp and Duboule, 2011)). However, a potential involvement of the regulatory topology, rather than the target end, remained to be assessed. In this study, by inverting the entire T-DOM, we could rule out the possibility that the physical order of various distant enhancers could play a major role in this mechanism, other than introducing a transcriptional delay, particularly visible in late-expressed *Hoxd* genes.

Finally, it is legitimate to wonder whether such moderate differences in the timing of gene activation could be detrimental to the development of the limb, considering that a close-to-normal expression pattern was resumed in E12.5 limb buds, except for the inversion of T-DOM containing the ectopic boundary where the anterior part of the domain remained absent even at later stages. This particular question was not addressed in this paper, since the detection of any loss of function phenotype would likely be hampered by the cooperative function of both the *HoxA* and *HoxD* clusters in developing limbs. Indeed, while their combined deletion led to very severe limb truncations, their deletion in isolation triggered much milder phenotypes (Kmita et al., 2005). While this functional complementation between these two gene clusters makes phenotypic analyses very complex in the mouse (it obliges to systematically remove the other gene cluster), it has allowed to study the underlying regulatory mechanisms in some details due to the persistence of a rather normal structure even after drastic chromosomal interventions.

In the above-mentioned alleles, it is thus difficult to anticipate whether or not any phenotype would be observable in the absence of the *HoxA* cluster. In the case of T-DOM inversion including the TAD border, it is however clear that the absence of transcripts at the anterior margin of the proximal expression domain would lead to an abnormal formation of the intermediate part of the limbs, as was shown in mice carrying a double inactivation of *Hoxa11* and *Hoxd11*, which displayed severely ill-formed forearms (Davis et al., 1995).

## MATERIALS AND METHODS

### Mouse strains

The *HoxD*^*del(CS38-40)*^ or *del(CS38-40)* allele was described in (Schep et al., 2016). The *HoxD*^*delCTCFs(CS38;CS40)*^ or *del(CTCFs), HoxD*^*inv(CS38-40)*^ or *inv(CS38-40), HoxD*^*inv(T-DOM)*^ or *inv(T-DOM)* and the *HoxD*^*inv(T-DOM)del(Bd)*^ or *inv(T-DOM)del(Bd)* alleles were generated through CRISPR/Cas9 editing technology using electroporation of mouse zygotes. The *del(CTCFs)* allele was derived from the *HoxD*^*delCTCF(CS38)*^ or *delCTCF(CS38)* allele and was also generated for this study using a gRNA directed against the consensus CTCF binding site located in region CS38, which generated a 26 bp large deletion. Subsequently, two gRNAs flanking region CS40 were designed to produce a 1’533 bp large deletion encompassing both CTCF binding sites at region CS40. For the *inv(CS38-40)* allele, two gRNAs were designed flanking the region CS38-40. Mice were genotyped either for a deletion or for an inversion of the region. Out of 48 specimens, only one mouse had an inversion, which we used as founder of the mutant line. For the *inv(T-DOM)* allele, a pair of gRNAs was directed at each end of the T-DOM regulatory domain. Out of 43 mice, only one had a full inversion, which was subsequently used to establish the mutant line. The breakpoints for the inversion were located 3’433 bp downstream of *Hoxd1* gene and 2’557 bp upstream of *Hnrnpa3* gene, inducing an inversion of 888’111 bp. This line was subsequently used to generate the *inv(T-DOM)del(Bd)* mice. For this latter allele, two gRNAs were designed flanking the boundary region now relocated in *inv(T-DOM)* close to *Hoxd1*, deleting a 17’303 bp large region. The *delCTCF(CS38)* and *inv(T-DOM)* alleles were generated after cloning the gRNAs into the pX330:hSpCas9 (Addgene ID 42230) vector and DNA injection into pronuclei. All other alleles were generated by electroporation of one-cell embryos with transcribed RNAs. gRNAs and genotyping primers are listed in Supplementary Table S1. All breakpoints were validated through Sanger sequencing and this information was used to generate the artificial mutant genomes which can be found in (http://doi.org/10.5281/zenodo.3826913) and a diagram in Fig S7.

### 4C-seq

Circular chromosome conformation capture (4C-seq) experiments were carried out as described in (Rodríguez-Carballo et al., 2017). Samples were micro-dissected from E12.5 or E9.5 forelimbs and placed in 10% FBS/PBS and incubated with collagenase at 37°C for 40 or 15 minutes respectively. Cell suspensions were then strained and fixed in 2% formaldehyde (FBS/PBS) for 10 minutes. For E12.5 experiments, between 10 and 12 pairs of distal or proximal forelimbs were used, while between 90 and 150 pairs of forelimbs were dissected for the E9.5 experiments. All E9.5 and the E12.5 *del(CTCFs)* 4C-seqs were conducted in embryos obtained from homozygous crosses, while all others were obtained from heterozygous crosses. The fastq from 4C-seq were demultiplexed, mapped and analysed using a local version of the pipeline that was present in HTSstation (http://htsstation.epfl.ch) (David et al., 2014) on the wild-type mm10 mouse genome (Fig. 2, 3, 4A, S2D, S2E) or on the *inv(T-DOM)* mm10 mouse genome (Fig. 4B, 5C, S4).The profiles were smoothened using a window size of 11 fragments. The track profiles were obtained using pygenometracks (Ramírez et al., 2018). The distribution of all 4C-seq viewpoints used in this work are listed in Supplementary Table S2.

### 4Cin

To compose the virtual Hi-C matrices we used the 4Cin package (Irastorza-Azcarate et al., 2018) and added an additional step to display the matrices with the linear distances. First, the profiles were reprocessed in four different ways to reduce the inherent variability of this computational approach. To this aim, the data values located at 0, <1, < 2 or <3 kb from the viewpoint were removed, then the data were smoothed using a window size of 11 fragments. Only data corresponding to chr2:74,400,000-75,800,000 (mm10) on the wild-type genome were used as input for the 4Cin.py script package (Irastorza-Azcarate et al., 2018) with default parameters. In order to obtain the best fitted representations, twenty different models generated by the 4Cin package were taken into account (five times for each of the four different exclusion pre-processings). They were clustered according to the Spearman correlation established between them. A Ward clustering was then applied using Euclidean distances (Fig. S8). The number of clusters was set so that the mean correlation of grouped models was <0.95 between clusters. For each cluster, an average was performed out of all the models and shown as supplementary figures when the cluster included more than one single model. In the main figures, an average of all twenty models was shown to better integrate all possible models, only if the clusters included more than one model. For example, for the *inv(CS38-40)* allele, only cluster 1 was considered as a bona fide representation, encompassing seventeen models, while the other three independent clusters were considered as outliers. In order to display the linear distances of the matrices, the virtual Hi-C output text file was converted to a cool file using a custom python script (https://github.com/lldelisle/scriptsForRodriguezCarballoEtAl2020.git). The track profiles were obtained using pygenometracks (Ramírez et al., 2018). The replicates and viewpoints used for each set of 4Cin are listed in Supplementary Tables S3 and S4.

### RNA extraction, RNA-seq and qPCR

Limb tissues were dissected and placed in individualized tubes containing RNAlater (Qiagen) and were frozen until genotyping and further processing. For E9.5 samples, once the limb buds were dissected, the rest of the embryo was fixed in 4% paraformaldehyde/PBS and stained with DAPI (Qiagen) for easy visualization of the somites under a microscope and characterization of the embryonic stage. All samples were processed following the RNeasy Microkit (Qiagen). RNA-seq libraries (one replicate per time-point) were generated from 100 ng of total RNA following the TruSeq Stranded mRNA protocol and sequenced on a HiSeq 2500 machine (100 bp single read). The gtf file used for STAR was based on Ensembl version 92 annotations (http://doi.org/10.5281/zenodo.3827120). Adapters were removed using cutadapt (v1.16; options -a GATCGGAAGAGCACACGTCTGAACTCCAGTCAC -m 15 –q 30) and aligned on mm10 using STAR version 2.6.0b-1 (Dobin et al., 2013) with ENCODE parameters. Only uniquely mapped reads were kept and coverage on each strand was obtained with bedtools genomecov v2.27.0. The track profiles were obtained using pygenometracks (Ramírez et al., 2018) and they show uniquely mapped reads normalized to the total number of uniquely mapped reads. For qPCR, RNA was retrotranscribed using the Promega GoScript Reverse Transcriptase (Promega). Custom SYBR probes were used for quantitative real-time PCR (qPCR) in a QuantStudio5 384-well block machine. All primers were described in (Delpretti et al., 2013; Montavon et al., 2008). For E9 samples, all values are relative to the average of the respective wild-type littermates. The number of replicates is mentioned in the figure legends. qPCR results were plotted using Graphpad Prism8.

### Chromatin Immunoprecipitation (ChIP)

Limb tissues were dissected and fixed in 1% formaldehyde/PBS for 10 min at room temperature, then incubated 3 min with Stop Solution from the ChIP-IT High Sensitivity Kit (Active Motif) and washed three times with PBS before being frozen at −80°C until further use. E9 wild-type limb tissues were pooled prior to fixation according to their somite stage, which was determined under a bright field microscope. Mutant samples were pooled according to their genotype prior to the experiment. Between 15 and 17 pairs of forelimb buds were used for each of the E9 ChIP experiments. For E12 ChIPs, the four entire limbs coming from one embryo were used in each experiment. All ChIP experiments were conducted following the ChIPmentation method (Schmidl et al., 2015) as in (Rodríguez-Carballo et al., 2019). Briefly, samples were Polytron minced and homogenized by douncing in Prep Buffer (ChIP-IT High Sensitivity Kit, Active Motif). They were then sonicated in 100 µl of sonication buffer (0.1%SDS, 50mM Tri-HCl pH8, 10mM EDTA ph8 and proteinase inhibitors) in a Bioruptor Pico sonicating device (Diagenode). All ChIPs were incubated overnight with the respective antibodies (CTCF Active Motif 61311; H3K27me3 Millipore 17-662; H3K27ac Diagenode C15410196; RAD21 abcam ab992) and precipitated after a 2h long incubation with magnetic beads (Dynabeads Protein A, Invitrogen 10001D). Washes were carried out in RIPA-LS, RIPA-HS and RIPA-LiCl. Beads were then resuspended in tagmentation buffer and incubated at 37°C for either 2 minutes (CTCF, RAD21, H3K27me3), or 10 minutes (H3K27ac) with 1 µl of Tn5 transposase (Illumina 15027865, from Nextera DNA Library Prep Kit 15028212). After washing with RIPA-LS and TE buffer, beads were incubated in elution buffer (10 mM Tris-HCl pH 8, 5mM EDTA pH 8, 300mM NaCl, 0.4% SDS) and proteinase K. DNA was then purified using Qiagen MiniElute kit and a qPCR was performed to determine the number of cycles to be applied during library amplification. DNA libraries were purified, size selected with CleanNGS magnetic beads (CleanNA) and sequenced in a HiSeq 4000 machine as 50 bp or 100 bp reads. The sequencing output was mapped and processed as in (Rodríguez-Carballo et al., 2019) without a normalization step. The track profiles were obtained using pygenometracks (Ramírez et al., 2018). In all figures, the orientations of the CTCF sites were obtained using the CTCFBSDB 2.0 database (http://insulatordb.uthsc.edu) based on the E9 CTCF ChIP data.

### Whole-mount in situ hybridization and LacZ transgenes

Whole-mount RNA in situ hybridization (WISH) was performed as in (Woltering et al., 2009) with the following modifications. E10.5 and E12.5 embryos were bleached in a 3% H_2_O_2_/PBS solution. After re-hydration, embryos were digested in a proteinase K solution (20µg/ml 10-12 min for E12.5 embryos, 10µg/ml 5 min for E10.5 embryos and 5µg/ml 4-5 minutes for E9.5 embryos). Digestion of E9.5 embryos was arrested by three quick washes in a 2mg/ml glycine solution, while E10.5 and E12.5 proteinase K digestions were stopped with a 10 min incubation in an acetic anhydride / triethanolamine solution. After mRNA probe hybridization and anti-DIG incubation, E9.5 embryos were washed several times overnight in maleic acid buffer, while E10.5 and E12.5 embryos’ washes were extended for a day.

The ELCR2 and ELCR3 regions were amplified and cloned into the betaGlobin reporter plasmid as in (Guerreiro et al., 2016) to generate the corresponding *LacZ* transgenes. These enhancer regions were amplified using the primer sequences GATGCTTGGCCTTAGCTCCT (Fw) and CTGTGGAAACGGAGCCAGAA (Rv) for ELCR2:*LacZ* and TCTCTGCCCATTCACTCTCATCA (Fw) and TTTTCTGTGCAGTGGCTGTGAC (Rv) for ELCR3:*LacZ*. The CS39:*lacZ* and CS65:*LacZ* transgenic lines were previously described (Beccari et al., 2016). All images were taken with a Leica MZFLIII and Leica M205 FA microscopes. A list with the number of replicates of each WISH is shown in Supplementary Table S5.

### Animal experimentation and ethics approval

All experiments were performed in agreement with the Swiss law on animal protection (LPA), under license number GE81/14 (to DD).

## Data availability

All sequencing data are deposited in the GEO database and can be found under the accession number GSE154189. All scripts used to generate the final outputs of Hi-C and 4C-seq (including figures) are available in https://github.com/lldelisle/scriptsForRodriguezCarballoEtAl2020.git.

## Competing interests

The authors declare that they have no competing interests.

### Acknowledgements

We want to thank Thi Hanh Nguyen Huynh, Marie-Laure Gadolini and Julien Codourey for their help with mice breeding, genotyping and transgenesis, as well as Mylène Docquier, Didier Chollet, Brice Petit and Christelle Barraclough from the Genomics platform at the University of Geneva. We also thank Ibai Irastorza-Azcarate and Asier Ullate-Agote for their help in implementing the 4Cin pipeline. We thank Aurélie Hintermann and other members of the Duboule laboratories for helpful discussions and suggestions.

## Author Contributions

ERC: project design, experimental work, manuscript writing.

LLD: bioinformatic analysis, amended the manuscript.

AW: experimental work (4C-seq), amended the manuscript.

LB: experimental work (transgenic design, conservation analysis).

SG, BM: pronuclear injections and embryo electroporation.

DD: project design, project funding, manuscript writing.

---

**Fig. S1.**
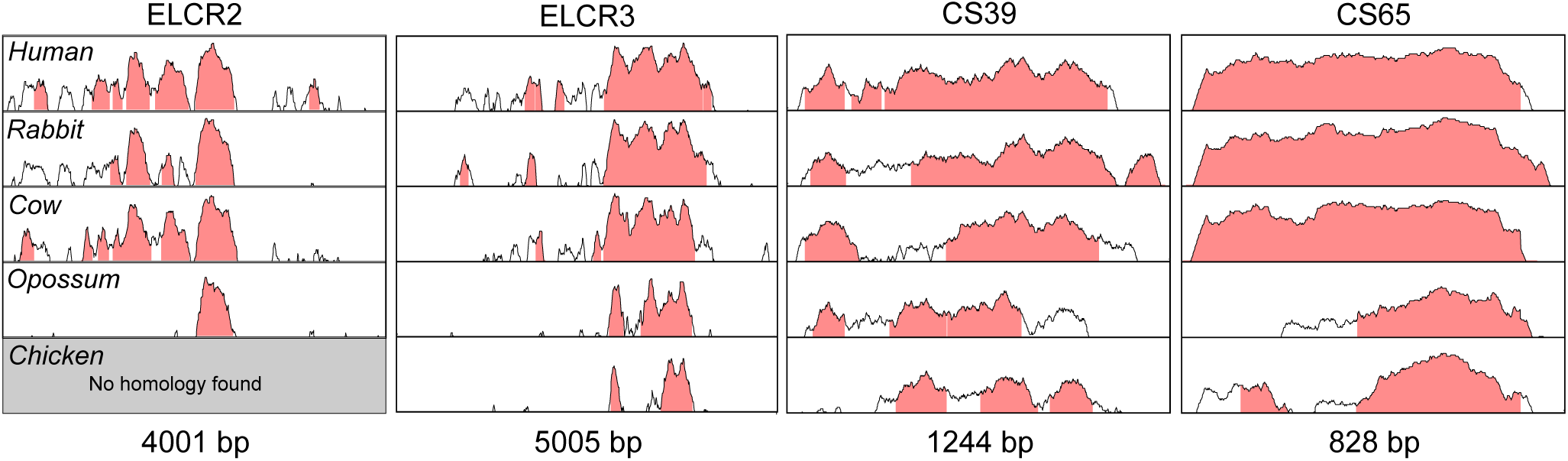
VISTA alignments of the enhancer sequences used in transgenic assays. Conservation was defined by at least 70% sequence identity over 100 bp. Various species were compared with the mouse sequences used to generate the transgenes (sizes are shown at the bottom). The y-axis ranges from 50 to 100% of identity.

**Fig. S2.**
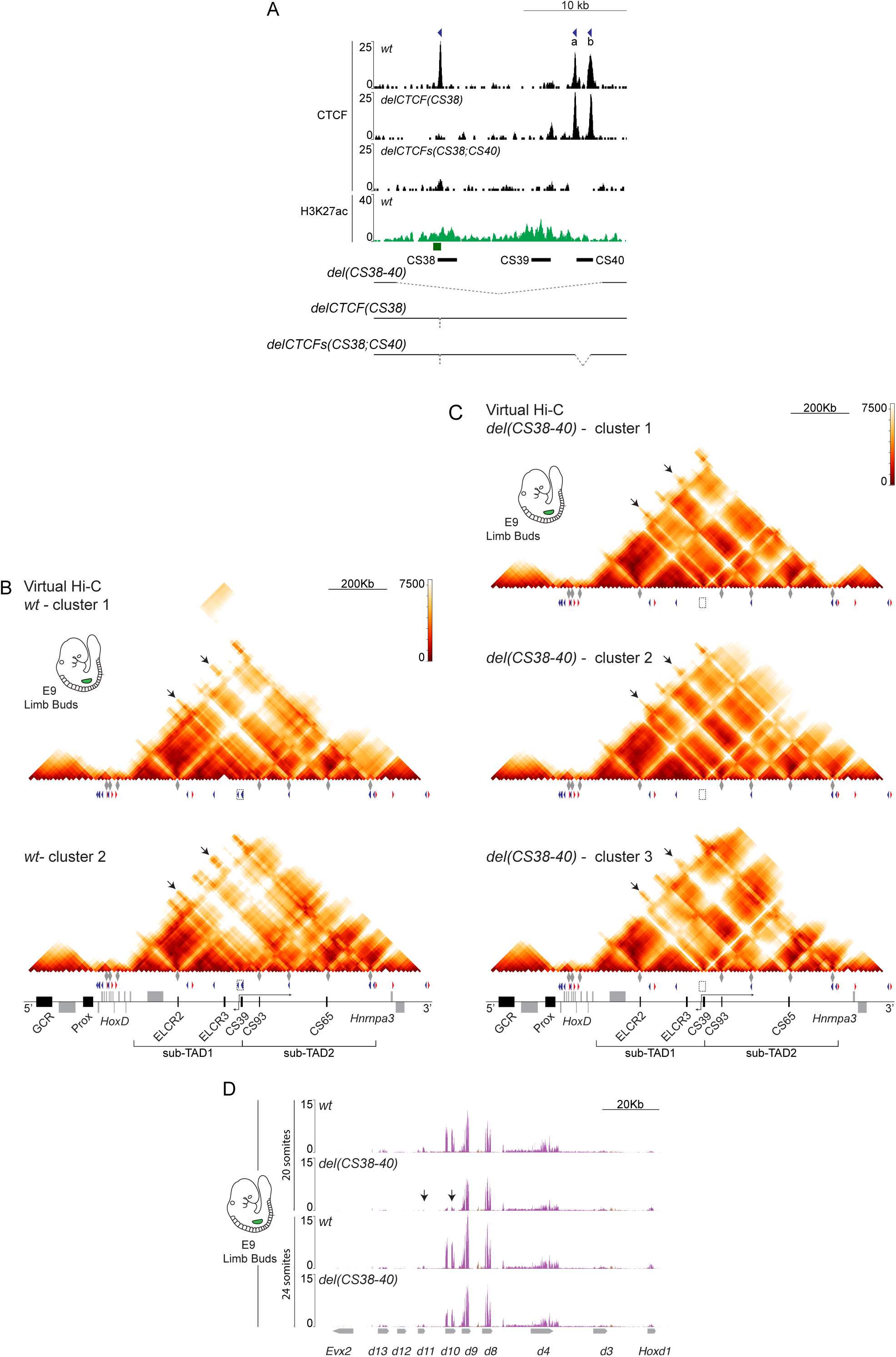
Topological importance of region CS38-40. **A**. On top are three tracks showing ChIP profiles of CTCF (black) in either wild-type, the *delCTCF(CS38)* or the *delCTCFs(CS38;CS40)* mutant alleles in E12.5 limb buds; below, the H3K27ac profile in wild-type E9 forelimb buds (green). The green box depicts the position of a CpG island, which grossly corresponds with the TSS of *Hog* and *Tog* lncRNAs. Bottom, schemes of the mutant lines associated with the deletions within region CS38-40. **B, C**. Matrices of contacts showing the two or three different profiles that could be obtained after clustering twenty different iterations with the 4Cin software of wild-type and *del(CS38-40)* mutant E9.5 limb buds (see Fig. S8). The arrows show the same regions as in Fig. 2. **D**. RNA-seq profiles of the *HoxD* region either in wild-type or in the mutant *del(CS38-40)* allele in E9.5 limb buds at somite stage 20 and 24. Arrows point to the delayed expression of *Hoxd10* and *Hoxd11*.

**Fig. S3.**
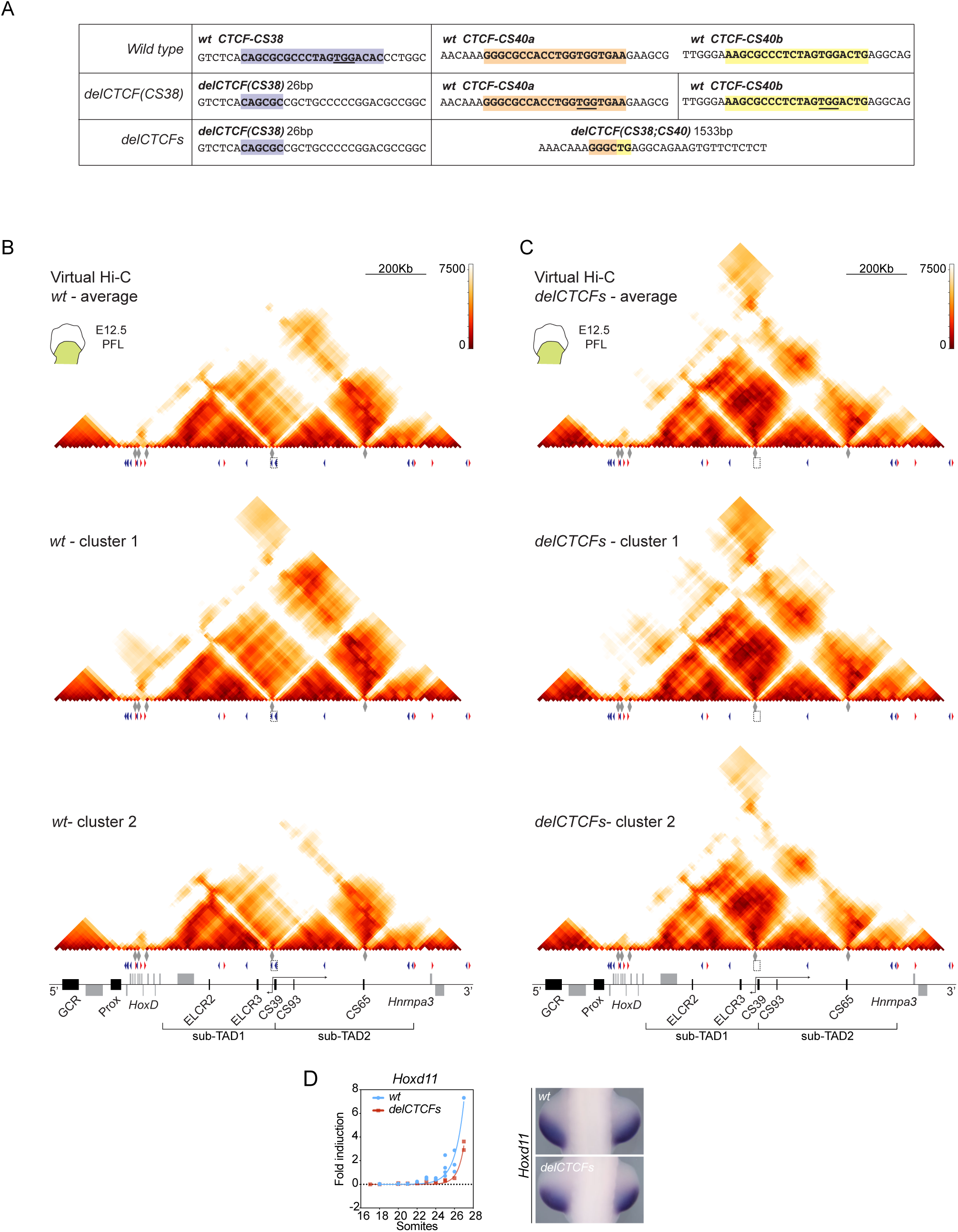
Deletion of CTCF binding sites in region CS38-40. **A**.DNA sequences of the CTCF binding sites in regions CS38 and CS40. The coloured boxes indicate the CTCF binding motives, also highlighted in bold. PAM sequences are underlined. **B, C**. 4Cin of E12.5 proximal limb buds dissected from either wild-type, or the *delCTCFs(CS38;CS40)* mutant stock. For each line, the virtual Hi-C on top shows the average of the two possible virtual Hi-Cs obtained if clusters were considered independently (see Fig. S8). Below each profile, the position of 4C-seq viewpoints and the orientated CTCF sites are shown as grey diamonds and red/blue arrowheads, respectively. **D**. Comparison of RT-qPCR values of *Hoxd11* in wild-type (n=22) and *delCTCFs(CS38;CS40)* (n=16) mutant limb buds along day E9. The WISH analysis of *Hoxd11* were carried out in E9.5 wild-type and mutant *delCTCFs(CS38;CS40)* specimens (approximately 29 somites).

**Fig. S4.**
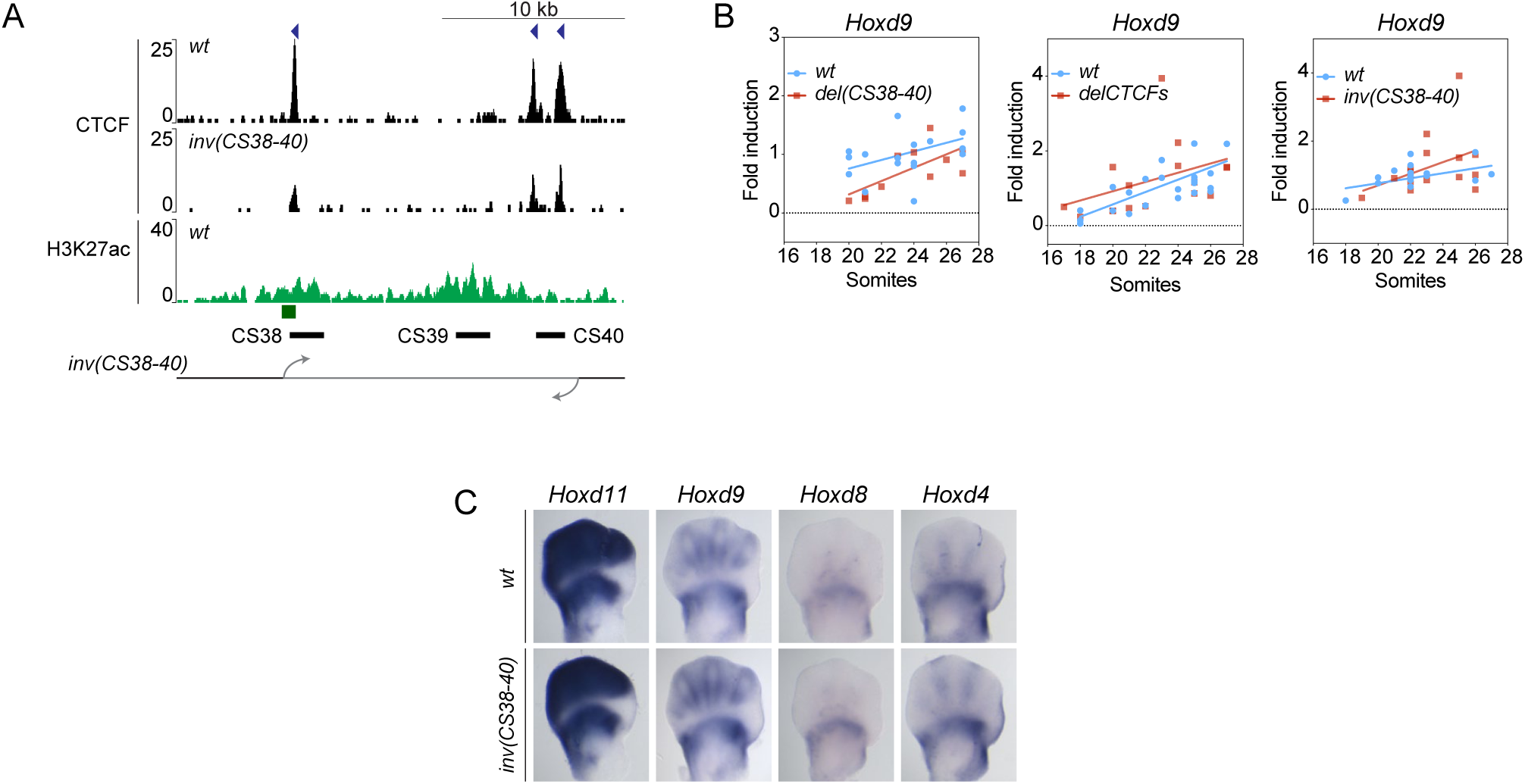
*Hoxd* genes expression after inversion of the sub-TAD border. **A**. On top are shown CTCF ChIP profiles (black) for both wild-type and the *inv(CS38-40)* mutant allele in E12.5 limb buds. Below is a H3K27ac profile (green) from E9 wild-type limb bud cells. The bottom track shows a scheme of the *inv(CS38-40)* mutant chromosome. **B**. Comparison of *Hoxd9* individual qPCR values along day E9. The plotted values correspond to the *del(CS38-40)* (n=11), *delCTCF(CS38;CS40)* (n=16) and *inv(CS38-40)* (n=16) alleles, compared to their respective wild-type littermates (n=17, 22 and 16, respectively). A regular linear fit is represented out of the real RT-qPCR values on each graph. **C**. WISH of *Hoxd11, Hoxd9, Hoxd8* and *Hoxd4* in either E12.5 wild-type, or *inv(CS38;CS40)* mutant forelimb buds.

**Fig. S5.**
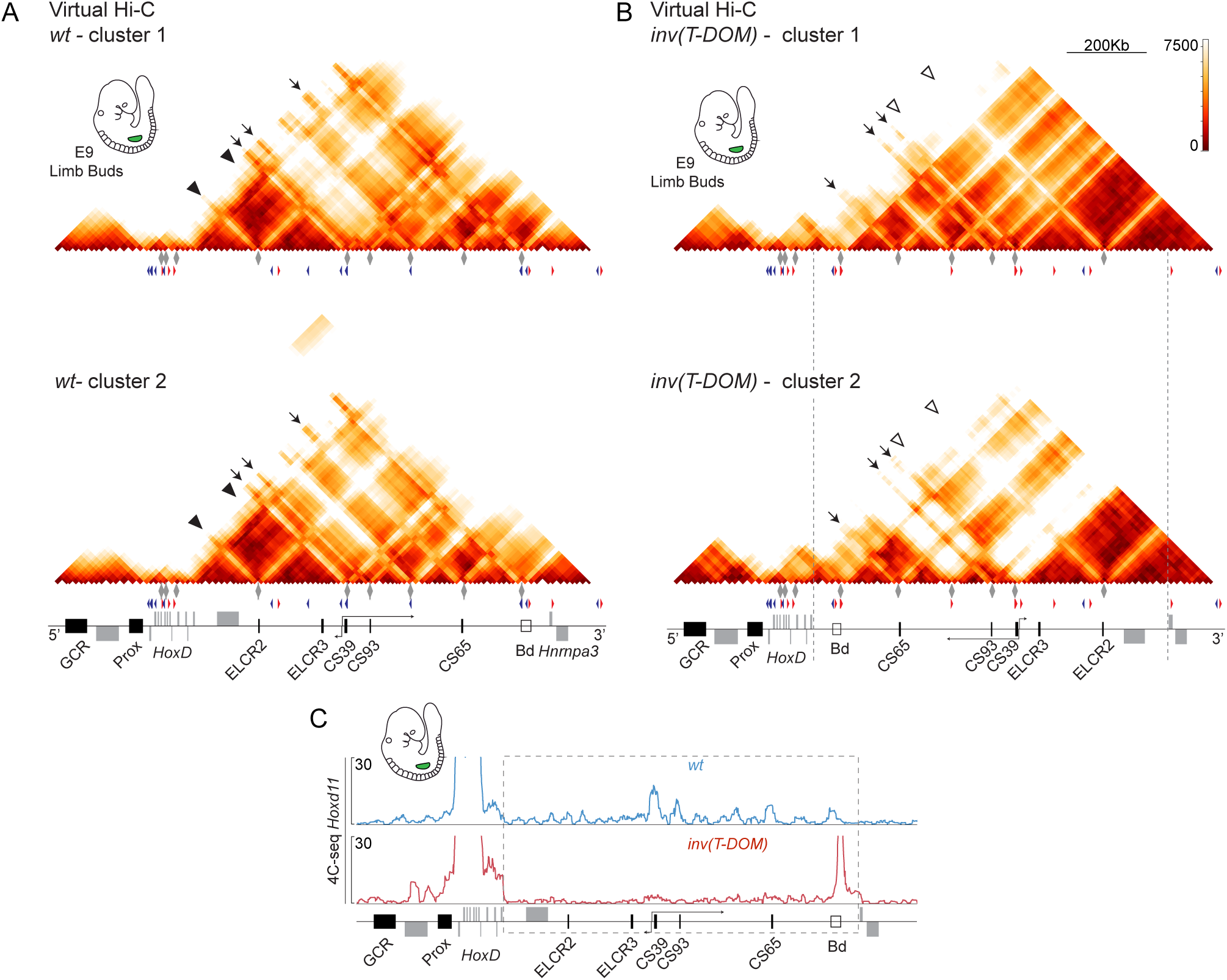
Modification of interactions in the *inv(T-DOM)* mutant allele. **A, B**. Matrices of contacts showing the two different profiles that could be obtained after clustering twenty different iterations with the 4Cin software of wild-type and *inv(T-DOM)* mutant E9.5 limb buds (see Fig. S8). Arrows and arrowheads show the same regions as in Fig.4. **C**. 4C-seq interaction profiles using *Hoxd11* as a bait in either wild-type (blue, top) or mutant *inv(T-DOM)* (red, bottom) cells from E9 forelimb buds. Both profiles were mapped on the wild-type genome and the inverted region is shown as a dashed grey box.

**Fig. S6.**
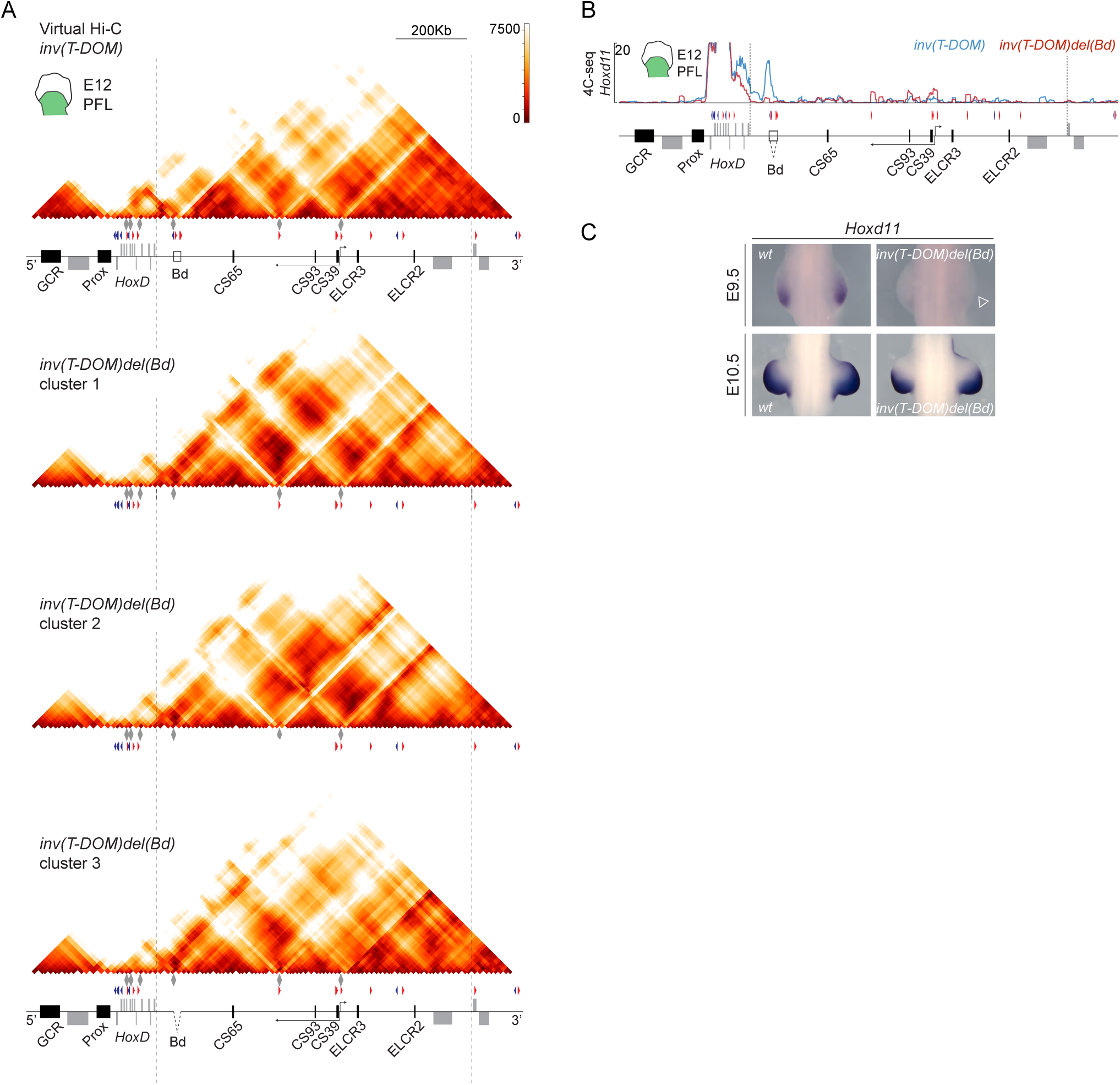
Modification of interactions in the *inv(T-DOM)del(Bd)* mutant allele. **A**. On top is a representation of the single virtual Hi-C profile obtained after twenty iterations of *inv(T-DOM)* mutant E12 limb data (same as in Fig. 5C). Below are representation of the three different clusters obtained from the mutant *inv(T-DOM)del(Bd)* data (see Fig. S8). The vertical dashed lines indicate the inverted region for both mutant lines. **B**. 4C-seq profiles using *Hoxd11* as a bait in either mutant *inv(T-DOM)* (blue) or mutant *inv(T-DOM)del(Bd)* (red) E12 proximal limb bud cells. Both profiles were mapped on the reconstructed *inv(T-DOM)* genome. **C**. WISH of *Hoxd11* using E9.5 (approximately 20 somites) and E10.5 wild-type and *inv(T-DOM)del(Bd)* mutant forelimb buds. The empty arrowhead indicates an absence of expression.

**Fig. S7.**
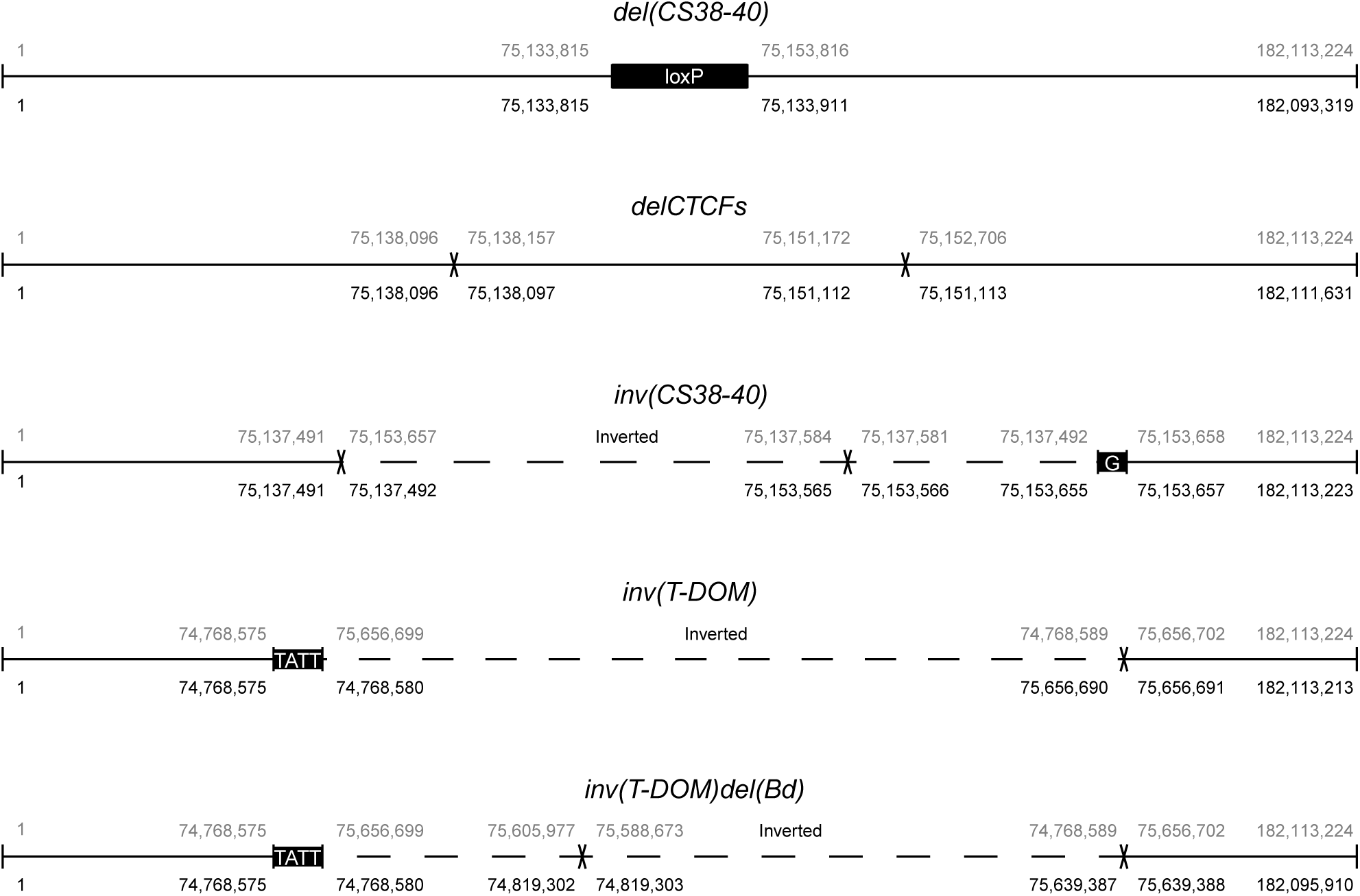
Scheme of the different mutant mouse stocks. The scheme shows the correspondence between the wild-type coordinates (on top, grey) and the new mutant coordinates (bottom, black) for each mutant line. The black boxes illustrate exogenous sequences such as LoxP (see *del(CS38-40)*) or randomly incorporated (see a ‘G’ in *inv(CS38-40)* and ‘TATT’ in *inv(T-DOM)* and *inv(T-DOM)del(Bd)*). Crosses indicate small deletions and dashed lines are for inverted regions. Note that the horizontal line represents the full extension of chromosome 2 and it is not on scale.

**Fig. S8.**
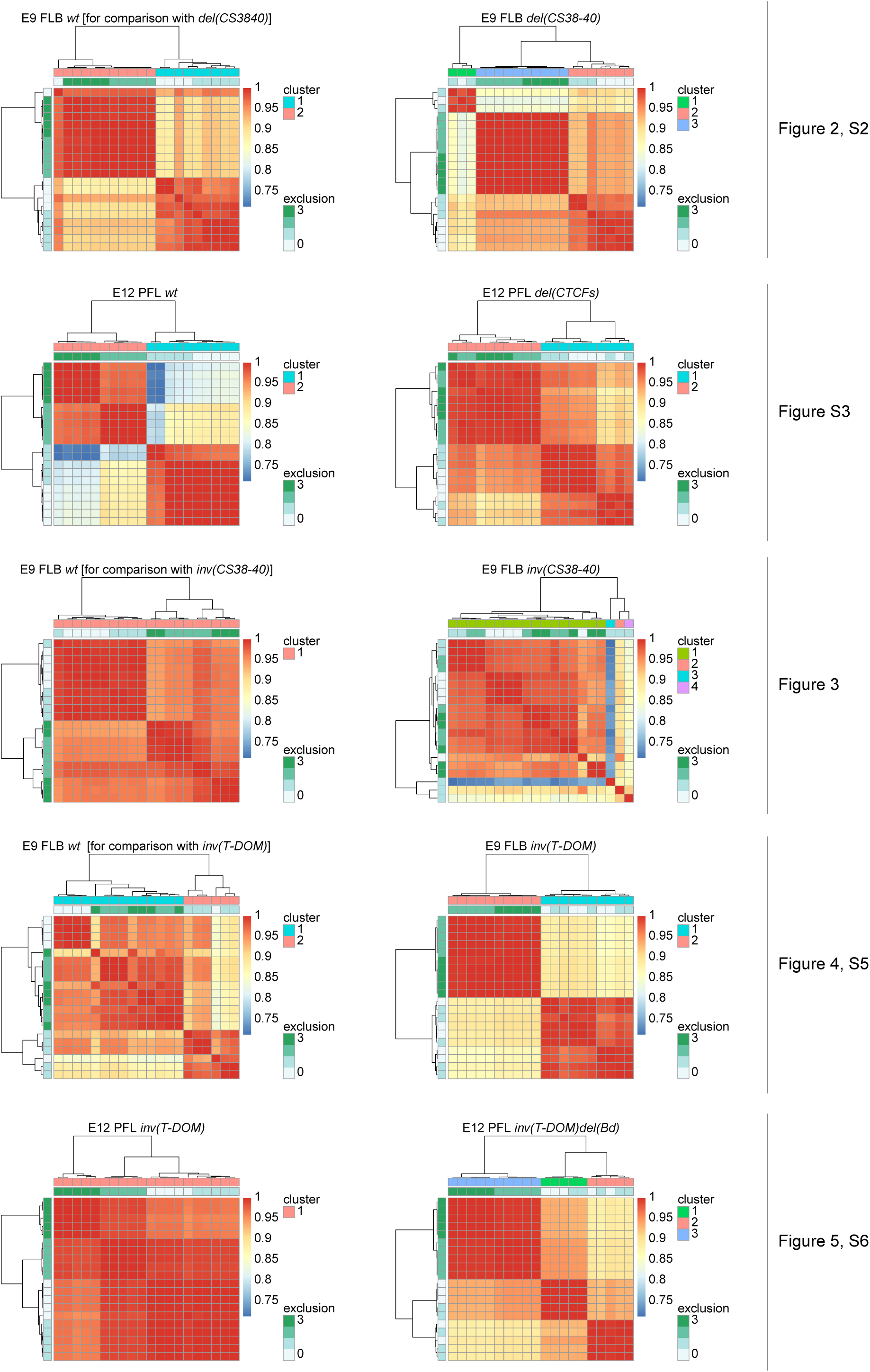
Clustering of 4Cin models. Heatmaps of Spearman correlation coefficient (blue to red scale bar) of twenty 4Cin iterations generated for each experiment. On the right of each heatmap, a green colored scale defines the exclusion size in the pre-processing for each model (see materials and methods). A ward clustering was applied using Euclidean distances on the correlation. Each experiment generated one to four clusters (see color code on the right of each diagram).

**Table S1.**
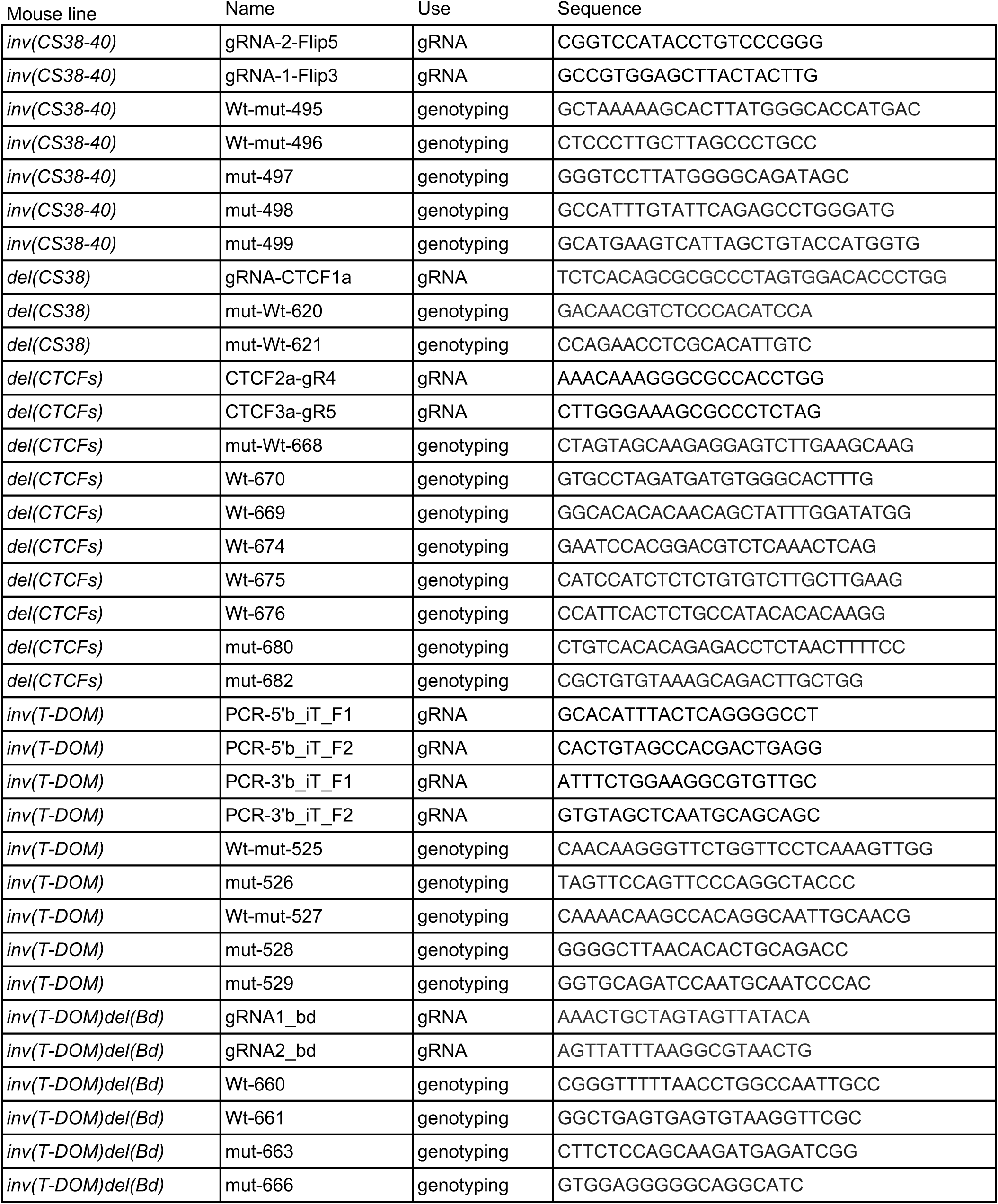
List of oligonucleotides. Sequences of oligonucleotides used as gRNA to produce the mutant lines generated for this study, as well as of the primers used to genotype them.

**Table S2.**
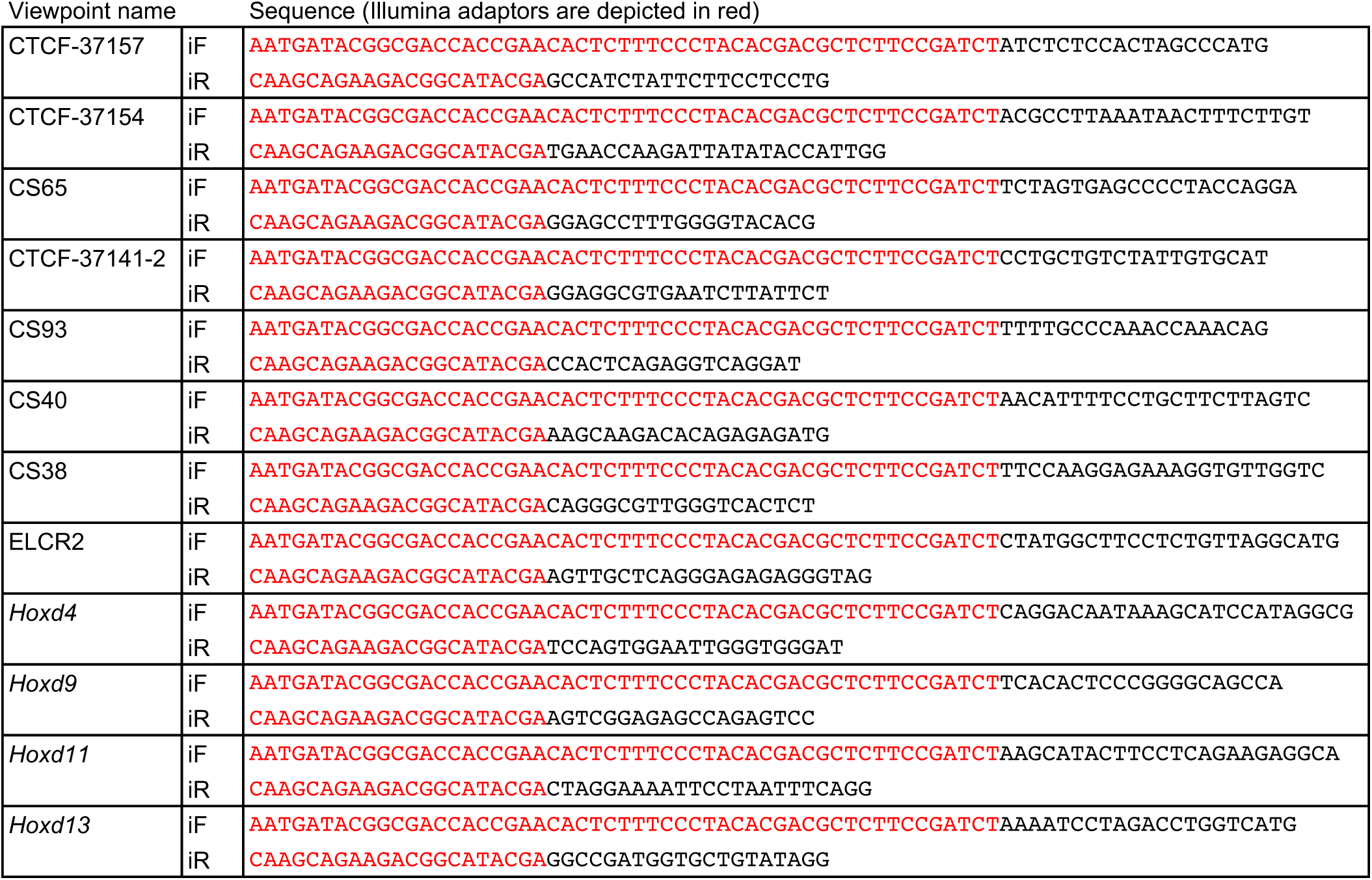
4C primers. Sequences of all the primers used as viewpoints in all 4C-seq experiments.

**Table S3.** 4C-seq experiments. Distribution of the replicates processed for each E9.5 4C-seq viewpoint.

Rodriguez-Carballo 2020 Supplementary Table S3
| Figure | viewpoint name | Mouse line | Embryonic Stage | Replicates | Source |
| --- | --- | --- | --- | --- | --- |
| Fig2, 3, 4 | CTCF-37154 | *Wt* | E9 | 1 | This study |
| Fig2, 3, 4 | CS65 | *Wt* | E9 | 1 | This study |
| Fig2, 3, 4 | CTCF-37141 | *Wt* | E9 | 2 | This study |
| Fig2, 3, 4 | CS93 | *Wt* | E9 | 1 | This study |
| Fig3, 4 | Cs40 | *Wt* | E9 | 1 | This study |
| Fig2, 3, 4 | ELCR2 | *Wt* | E9 | 1 | This study |
| Fig2, 3, 4 | *Hoxd4* | *Wt* | E9 | 1 | This study |
| Fig2, 3, 4 | *Hoxd9* | *Wt* | E9 | 1 | This study |
| Fig2, 3, 4 | *Hoxd11* | *Wt* | E9 | 1 | This study |
| Fig3 | *Hoxd13* | *Wt* | E9 | 1 | This study |
| Fig2 | CTCF-37154 | *del(CS38-40)* | E9 | 1 | This study |
| Fig2 | CS65 | *del(CS38-40)* | E9 | 2 | This study |
| Fig2 | CTCF-37141 | *del(CS38-40)* | E9 | 2 | This study |
| Fig2 | CS93 | *del(CS38-40)* | E9 | 1 | This study |
| Fig2 | ELCR2 | *del(CS38-40)* | E9 | 2 | This study |
| Fig2 | *Hoxd4* | *del(CS38-40)* | E9 | 2 | This study |
| Fig2 | *Hoxd9* | *del(CS38-40)* | E9 | 2 | This study |
| Fig2 | *Hoxd11* | *del(CS38-40)* | E9 | 2 | This study |
| Fig3 | CTCF-37154 | *inv(CS38-40)* | E9 | 1 | This study |
| Fig3 | CS65 | *inv(CS38-40)* | E9 | 1 | This study |
| Fig3 | CTCF-37141 | *inv(CS38-40)* | E9 | 1 | This study |
| Fig3 | CS93 | *inv(CS38-40)* | E9 | 1 | This study |
| Fig3 | Cs40 | *inv(CS38-40)* | E9 | 2 | This study |
| Fig3 | ELCR2 | *inv(CS38-40)* | E9 | 2 | This study |
| Fig3 | *Hoxd4* | *inv(CS38-40)* | E9 | 2 | This study |
| Fig3 | *Hoxd9* | *inv(CS38-40)* | E9 | 1 | This study |
| Fig3 | *Hoxd11* | *inv(CS38-40)* | E9 | 2 | This study |
| Fig3 | *Hoxd13* | *inv(CS38-40)* | E9 | 1 | This study |
| Fig4 | CTCF-37154 | *inv(T-DOM)* | E9 | 1 | This study |
| Fig4 | CS65 | *inv(T-DOM)* | E9 | 1 | This study |
| Fig4 | CTCF-37141 | *inv(T-DOM)* | E9 | 1 | This study |
| Fig4 | CS93 | *inv(T-DOM)* | E9 | 1 | This study |
| Fig4 | Cs40 | *inv(T-DOM)* | E9 | 1 | This study |
| Fig4 | ELCR2 | *inv(T-DOM)* | E9 | 1 | This study |
| Fig4 | *Hoxd4* | *inv(T-DOM)* | E9 | 1 | This study |
| Fig4 | *Hoxd9* | *inv(T-DOM)* | E9 | 1 | This study |
| Fig4 | *Hoxd11* | *inv(T-DOM)* | E9 | 1 | This study |

**Table S4.** 4C-seq experiments. Distribution of the replicates processed for each E12.5 4C-seq viewpoint.

Rodriguez-Carballo 2020 Supplementary Table S4
| Figure | Viewpoint name | Mouse line | Embryonic Stage | Replicates | Source |
| --- | --- | --- | --- | --- | --- |
| Fig5 | CTCF-37157 | *inv(T-DOM)del(Bd)* | E12 | 1 | This study |
| Fig5 | CTCF-37141 | *inv(T-DOM)del(Bd)* | E12 | 1 | This study |
| Fig5 | CS38 | *inv(T-DOM)del(Bd)* | E12 | 1 | This study |
| Fig5 | *Hoxd4* | *inv(T-DOM)del(Bd)* | E12 | 1 | This study |
| Fig5 | *Hoxd9* | *inv(T-DOM)del(Bd)* | E12 | 1 | This study |
| Fig5 | *Hoxd11* | *inv(T-DOM)del(Bd)* | E12 | 1 | This study |
| Fig5 | CTCF-37157 | *inv(T-DOM)* | E12 | 1 | This study |
| Fig5 | CTCF-37141 | *inv(T-DOM)* | E12 | 1 | This study |
| Fig5 | CS38 | *inv(T-DOM)* | E12 | 1 | This study |
| Fig5 | *Hoxd4* | *inv(T-DOM)* | E12 | 1 | This study |
| Fig5 | *Hoxd9* | *inv(T-DOM)* | E12 | 1 | This study |
| Fig5 | *Hoxd11* | *inv(T-DOM)* | E12 | 1 | This study |
| FigS2 | CS65 | *Wt* | E12 | 2 | This study and GEO GSM2713698 |
| FigS2 | CS38 | *Wt* | E12 | 6 | This study and GEO GSM2713680 |
| FigS2 | *Hoxd4* | *Wt* | E12 | 5 | This study and GEO GSM2713672 |
| FigS2 | *Hoxd9* | *Wt* | E12 | 1 | This study |
| FigS2 | *Hoxd11* | *Wt* | E12 | 3 | This study |
| FigS2 | CS65 | *del(CTCFs)* | E12 | 1 | This study |
| FigS2 | CS38 | *del(CTCFs)* | E12 | 1 | This study |
| FigS2 | *Hoxd4* | *del(CTCFs)* | E12 | 1 | This study |
| FigS2 | *Hoxd9* | *del(CTCFs)* | E12 | 1 | This study |
| FigS2 | *Hoxd11* | *del(CTCFs)* | E12 | 1 | This study |

**Table S5.** WISH replicates. Distribution of the replicates processed for each WISH shown in this article.

Rodriguez-Carballo 2020 Supplementary Table S5
| Figure | Experiment | Mouse line | Embryonic day | Replicates |
| --- | --- | --- | --- | --- |
| Fig2 | *Hoxd11* WISH | *Wt* | E9.5 | 3 |
| Fig2 | *Hoxd11* WISH | *del(CS38-40)* | E9.5 | 3 |
| Fig3 | *Hoxd11* WISH | *Wt* | E9.5 | 4 |
| Fig3 | *Hoxd11* WISH | *inv(CS38-40)* | E9.5 | 2 |
| Fig4 | *Hoxd11* WISH | *Wt* | E9.5 | 2 |
| Fig4 | *Hoxd11* WISH | *inv(T-DOM)* | E9.5 | 5 |
| Fig4 | *Hoxd11* WISH | *Wt* | E10.5 | 2 |
| Fig4 | *Hoxd11* WISH | *inv(T-DOM)* | E10.5 | 3 |
| Fig5 | *Hoxd11* WISH | *Wt* | E12.5 | 1 |
| Fig5 | *Hoxd11* WISH | *inv(T-DOM)* | E12.5 | 2 |
| Fig5 | *Hoxd11* WISH | *Wt* | E12.5 | 1 |
| Fig5 | *Hoxd11* WISH | *inv(T-DOM)del(Bd)* | E12.5 | 1 |
| Fig5 | *Hoxd9* WISH | *Wt* | E12.5 | 1 |
| Fig5 | *Hoxd9* WISH | *inv(T-DOM)* | E12.5 | 2 |
| Fig5 | *Hoxd9* WISH | *Wt* | E12.5 | 2 |
| Fig5 | *Hoxd9* WISH | *inv(T-DOM)del(Bd)* | E12.5 | 1 |
| FigS2 | *Hoxd11* WISH | *Wt* | E9.5 | 8 |
| FigS2 | *Hoxd11* WISH | *del(CTCFs)* | E9.5 | 4 |
| FigS3 | *Hoxd4* WISH | *Wt* | E12.5 | 1 |
| FigS3 | *Hoxd4* WISH | *inv(CS38-40)* | E12.5 | 1 |
| FigS3 | *Hoxd8* WISH | *Wt* | E12.5 | 1 |
| FigS3 | *Hoxd8* WISH | *inv(CS38-40)* | E12.5 | 2 |
| FigS3 | *Hoxd9* WISH | *Wt* | E12.5 | 3 |
| FigS3 | *Hoxd9* WISH | *inv(CS38-40)* | E12.5 | 1 |
| FigS3 | *Hoxd11* WISH | *Wt* | E12.5 | 1 |
| FigS3 | *Hoxd11* WISH | *inv(CS38-40)* | E12.5 | 2 |
| FigS5 | *Hoxd11* WISH | *Wt* | E9.5 | 2 |
| FigS5 | *Hoxd11* WISH | *inv(T-DOM)del(Bd)* | E9.5 | 2 |
| FigS5 | *Hoxd11* WISH | *Wt* | E10.5 | 1 |
| FigS5 | *Hoxd11* WISH | *inv(T-DOM)del(Bd)* | E10.5 | 1 |

